# Brunner syndrome associated MAOA dysfunction in human dopaminergic neurons results in NMDAR hyperfunction and increased network activity

**DOI:** 10.1101/2020.10.28.359224

**Authors:** Yan Shi, Jon-Ruben van Rhijn, Maren Bormann, Britt Mossink, Monica Frega, Hatice Recaioglu, Marina Hakobjan, Teun Klein Gunnewiek, Chantal Schoenmaker, Elizabeth Palmer, Laurence Faivre, Sarah Kittel-Schneider, Dirk Schubert, Han Brunner, Barbara Franke, Nael Nadif Kasri

## Abstract

**Background:** Monoamine neurotransmitter abundance affects motor control, emotion, and cognitive function and is regulated by monoamine oxidases. Amongst these, monoamine oxidase A (MAOA) catalyzes the degradation of dopamine, norepinephrine, and serotonin into their inactive metabolites. Loss-of-function mutations in the X-linked *MAOA* gene cause Brunner syndrome, which is characterized by various forms of impulsivity, maladaptive externalizing behavior, and mild intellectual disability. Impaired MAOA activity in individuals with Brunner syndrome results in bioamine aberration, but it is currently unknown how this affects neuronal function.

**Methods:** We generated human induced pluripotent stem cell (hiPSC)-derived dopaminergic (DA) neurons from three individuals with Brunner syndrome carrying different mutations, and used CRISPR/Cas9 mediated homologous recombination to rescue MAOA function. We used these lines to characterize morphological and functional properties of DA neuronal cultures at the single cell and neuronal network level *in vitro*.

**Results:** Brunner syndrome DA neurons showed reduced synaptic density but hyperactive network activity. Intrinsic functional properties and α-amino-3-hydroxy-5-methyl-4-isoxazolepropionic acid receptor (AMPAR)-mediated synaptic transmission were not affected by MAOA dysfunction. Instead, we show that the neuronal network hyperactivity is mediated by upregulation of the *GRIN2A* and *GRIN2B* subunits of the N-methyl-D-aspartate receptor (NMDAR), and rescue of *MAOA* results in normalization of NMDAR function as well as restoration of network activity.

**Conclusions:** Our data suggest that MAOA dysfunction in Brunner syndrome increases activity of dopaminergic neurons through upregulation of NMDAR function, which may contribute to Brunner syndrome associated phenotypes.

## Introduction

Dopamine, serotonin, and noradrenaline all belong to the class of monoamine neurotransmitters. They are prevalent throughout the brain, and their abundance influences brain development, function and behavior(1, 2). Monoamine neurotransmitter related activity is tightly regulated, and dysregulation of monoaminergic pathways is associated with several neuropsychiatric disorders including schizophrenia, major depressive disorder, autism spectrum disorder (ASD), and attention deficit/hyperactivity disorder (ADHD). Monoamine oxidases (MAOs) catabolize monoaminergic neurotransmitters(3) and thereby regulate the monoamine concentration in the brain. Disruption of MAO activity can have profound consequences on normal brain function(4). One disorder in which MAO function is strongly affected is Brunner syndrome, a neurodevelopmental disorder characterized by hemizygous mutations in the X-linked monoamine oxidase-A (*MAOA*) gene. Brunner syndrome was first described in large Dutch kindred with non-dysmorphic borderline intellectual disability (ID), and prominent impulsivity and maladaptive externalizing behavior(5, 6). More recently, three more families have been reported with Brunner syndrome, strengthening the link between MAOA dysfunction and Brunner syndrome. In all families, individuals carry either nonsense or missense mutations of *MAOA*(7, 8).

The monoaminergic system has been associated with regulation of aggressive behavior, both in wildtype animal models(9–12) and genetic models of neurodevelopmental disorders(13, 14). For example, hemizygous *Maoa* mutant mice show abnormally high levels of aggressive behavior and disturbed monoamine metabolism(15–17). Furthermore, *Maoa-deficient* mice display alterations in brain development, with aberrant organization of the primary somatosensory cortex(15) and increased dendritic arborization of pyramidal neurons in the orbitofrontal cortex(17). Postnatal reduction of serotonin levels in *Maoa*-deficient mice partially corrected some of these developmental abnormalities in the cortex(18). On the molecular level, *Maoa* has been implicated in the regulation of synaptic neurotransmitter receptors, as *Maoa* knockout mice show increased N-methyl-D-aspartate (NMDA) receptor subunit expression in the prefrontal cortex(19). Taken together, these data suggest that dysfunction of MAOA in rodents leads to both structural and functional alterations during brain development.

MAOA is expressed in different neuronal as well as glial cell types in the brain*(20)*. This complex interplay of multiple monoaminergic pathways in brain function creates a challenge in disentangling the cell-type specific roles of MAOA during neurodevelopment and in the regulation of normal brain activity. So far, particular emphasis has been given to the serotonergic system in MAOA research(21), as MAOA dysfunction results in increased serotonin levels in both humans and mice(5, 7, 8, 15, 17, 22). Indeed, dysfunction of the serotonergic system is associated with increased aggression and impulsivity(23). However, the expression pattern of MAOA suggests it is primarily expressed in catecholaminergic (dopaminergic and noradrenergic) neurons(21, 24), whereas expression in serotonergic neurons is variable and decreases during development in the mouse brain(25). Abundant expression of MAOA in dopaminergic (DA) neurons(24, 26) coincides with the finding that changes in dopaminergic neuronal activity also directly affect impulsive and aggressive behavior(27, 28).

Current advances in the generation of human induced pluripotent stem cell (hiPSC) induced neurons enable us to generate cultures of defined cell types, which provides us opportunities to disentangle the complexities that underlie interactions of multiple monoaminergic pathways. We generated hiPSC-derived DA neurons from healthy individuals and individuals with Brunner syndrome carrying missense or nonsense mutations to investigate the cellular and molecular mechanisms underlying MAOA dysfunction in a homogenous human DA neuronal network. Combining data on morphological analysis, gene expression, single-cell electrophysiology, and neuronal network activity using microelectrode arrays (MEAs), we show that increased network activity in MAOA-deficient DA neuronal networks is associated with increased expression of the N-Methyl-D-Aspartate receptor (NMDAR) subunits *GRIN2A* and *GRIN2B* and increased NMDAR function. Rescue of a *MAOA* missense mutation by CRISPR/Cas9 resulted in restoration of *GRIN2A* and *GRIN2B* expression, NMDAR function, and neuronal network activity to control values. Taken together, this work suggests that increased network activity in DA neurons from individuals with Brunner syndrome is causally linked to NMDAR hyperfunction.

## Methods and Materials

Methods and materials are described in greater detail in the Supplemental Methods and Materials.

### Cell Culture of hiPSCs and neuron differentiation

Control hiPSC lines were derived from dermal fibroblasts of male healthy volunteers(29),(30). The hiPSC lines ME2, ME8 and NE8 were derived from dermal fibroblast biopsies of male individuals diagnosed with Brunner syndrome described previously(5–8). Ethical approval for the study was obtained for all sites separately by local ethics committees. Written informed consent was given by the parents or legal representatives of the participants. All hiPSC reprogramming and characterization of the pluripotency markers was done by the Radboudumc Stem Cell Technology Center (SCTC) (**Supplementary Figure 1**). The hiPSCs colonies were split into single cells, and differentiated to a DA neuron identity using small molecules(31). Rat astrocytes (prepared as previously described(32)) were cocultured with DA neuron progenitors to promote maturation.

### Gene expression analysis and immunocytochemistry

RNA was isolated with the RNeasy Mini Kit (Qiagen) and retro-transcribed into cDNA by iScript cDNA Synthesis Kit (Bio-Rad Laboratories, Inc) according to the manufacturer’s instructions. We measured gene expression profiles using quantitative real-time PCR (qRT-PCR). Primers are listed in **Supplementary Table S1**. DA neurons plated on glass coverslips were used for immunocytochemistry and imaged using a Zeiss AxioImager Z1 with apotome (Carl Zeiss AG, Germany). Synapse density was assessed through manual counting using Fiji software.

### Neuronal reconstruction

Widefield fluorescent images of MAP2-labelled hiPSC-derived DA neurons were reconstructed using Neurolucida 360 (Version 2017.01.4, Microbrightfield Bioscience, Williston, USA). All the morphological data were acquired and analyzed blind to the genetic background of the neurons.

### MEA recording and Single-cell electrophysiology

DA neuron progenitors (day *in vitro* 20, DIV20) were plated on 6-Well or 24-well MEA devices (Multichannel Systems, MCS GmbH, Reutlingen, Germany). The neuronal network activity of DIV73 DA neurons was measured and analyzed as described(32–34). Single-cell electrophysiological recordings on coverslip containing DIV73 DA neurons were conducted under continuous perfusion with oxygenated (95% O2 / 5% CO2) and heated (32°C) recording artificial cerebrospinal fluid (ACSF).

### Statistical analysis

Statistical analysis of the data was performed with GraphPad PRISM (GraphPad Software, Inc, USA). Data is always shown as mean ± SEM. A detailed overview of all averaged data can be found in supplementary data tables S3-S7. Mann-Whitney U test, unpaired Student’s T test or one-way ANOVA with Dunnet’s correction for multiple-comparisons was used for statistical analysis. P<0.05 was considered significant.

## Results

### Generation of DA neurons derived from individuals with and without Brunner syndrome

We generated hiPSCs from two healthy subjects and three Brunner syndrome patients from independent families (**Figure 1a**, for extended information see **Supplementary Table S1**). ME2 and ME8 had a missense mutation in exon 2 (c.133C>T, p.R45W(8)) and exon 8 (c.797_798delinsTT, p.C266F(7)), respectively. These mutations are both located in the flavin adenine dinucleotide (FAD)-binding domain of MAOA(8) **(Figure 1b**). NE8 had a nonsense mutation in *MAOA* leading to a premature stop codon (c.886C>T, p.Q296**)(5). All patients were known to display elevated serotonin and disturbed monoamine metabolite levels in serum and urine(3, 6–8). All selected clones expressed the pluripotency markers OCT4, TRA-1-81, NANOG, and SSEA4 (**Supplementary Figure 1**), and *MAOA* mutations were confirmed by Sanger sequencing in the fibroblast-derived hiPSC lines from individuals with Brunner syndrome (**Figure 1a**).

**Figure 1.**
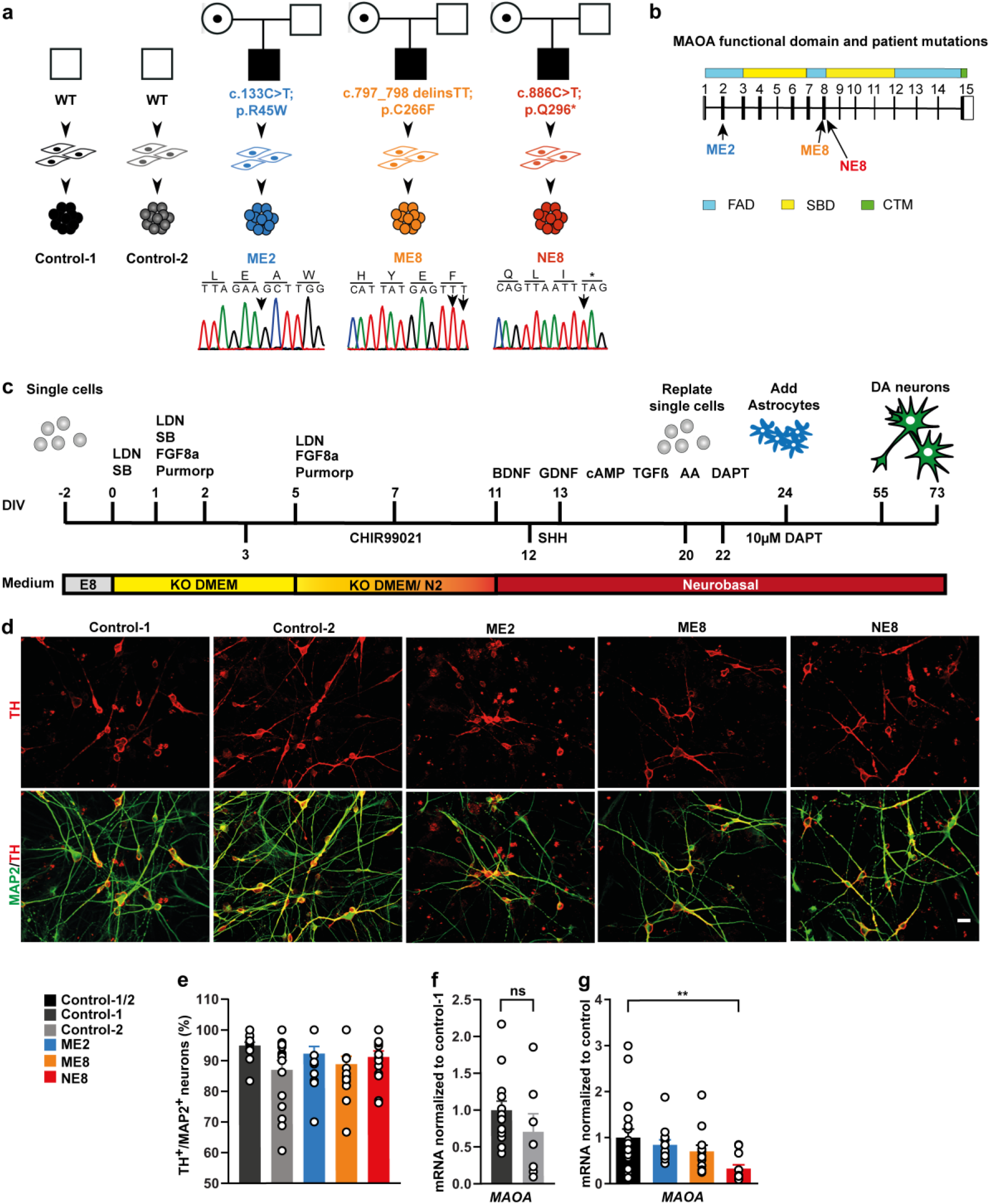
Differentiation of DA neurons derived from human induced pluripotent stem cells (hiPSCs). (a) Scheme of control and patient hiPSC lines used in the study. The monoamine oxidase A (*MAOA*) mutations were confirmed by sanger sequencing. (b) Location of the different mutations within the *MAOA* gene and protein domain. FAD (blue boxes), flavin adenine dinucleotide binding domains; SBD (yellow boxes), substrate-binding domain; CTM (green box), C-terminal membrane region. (c) Schematic overview of the protocol used to generate DA neurons from hiPSCs. (d) Representative images of DIV 55 DA neurons labeled by TH (red) and MAP2 (green) (Scale bar = 20 μm). (e) The percentage of TH-positive neurons (among MAP2-positive cells) at DIV 55. Sample size: Control-1 N=15, Control-2 N=16, ME2 N=15, ME8 N=15, NE8 N=15. (f) *MAOA* mRNA expression in control DIV 73 DA neurons. Sample size: Control-1 N=13, Control 2 N=7. (g) Comparison of *MAOA* mRNA expression between control and patient lines (Control vs NE8 *P=0.0076*). Sample size: ME2 N=12, ME8 N=12, NE8 N=12. All data represent means ± SEM. One-Way ANOVA with Dunnett’s correction for multiple testing was used to compare between patient lines and control lines. *\*\*P<0.01*.

We differentiated hiPSCs into a homogenous population of DA neurons using small molecules(31) (**Figure 1c**). For all experiments, DA neurons were co-cultured with rodent astrocytes to facilitate neuronal development and network maturation. Neuronal identity was confirmed by microtubule-associated protein 2 (MAP2) expression and DA neuron identity by expression of the dopaminergic neuron marker tyrosine hydroxylase (TH) after 55 days *in vitro* (DIV 55, **Figure 1d**). All hiPSC lines were able to differentiate into TH/MAP2 double-positive neurons at similar efficiency (**Figure 1e**).(31)*MAOA* mRNA levels were similar between control lines (**Figure 1f**), and the ME2 and ME8 lines (**Figure 1g**). As expected, mRNA levels of *MAOA* were reduced in the NE8 line compared to healthy controls (**Figure 1g**). This is likely caused by nonsense-mediated mRNA decay, which has been reported in human fibroblasts with the same mutation(35). Of note, ME2 and NE8 carry an allele of the variable number tandem repeat (VNTR) polymorphism in the MAOA promoter region associated with high gene expression (36, 37), whereas control-1, control-2 and ME8 carry an allele associated with low expression (**Supplementary Figure 2**). These alleles have previously been suggested to affect *MAOA* expression differentially using luciferase assays in immortalized cell lines(38). However, in hiPSC-derived DA neurons, *MAOA* expression does not seem to be affected by this polymorphism.

### MAOA dysfunction affects synapse density in DA neurons

It has been shown that dendritic arborization of pyramidal neurons in the orbitofrontal cortex is increased in *Maoa* hemizygous knockout mice(17). However, it is unclear whether this is a direct effect of impaired MAOA expression or function. We therefore immunostained MAP2 to identify the soma and dendrites of DA neurons (**Supplementary Figure 3**) and used quantitative morphometric analysis to assess whether MAOA dysfunction directly affects neuronal somatodendritic morphology (**Figure 2a**). Comparison of DA neurons from healthy individuals and those with Brunner syndrome revealed cell-line specific alterations of DA neuron morphology after 73 days of differentiation (DIV 73). ME8 DA neurons showed a significant increase in soma size (**Figure 2b**) and dendrite complexity including dendritic nodes, length and Sholl analysis (**Figure 2c, d, f,** respectively) compared to controls. Whilst NE8 DA neurons showed no differences from controls in dendritic complexity (**Figure 2c-f**). Both ME8 and ME2 DA neurons showed a significant increase in the total dendritic span (Convex Hull analysis, **Figure 2g**), which suggests that missense mutations in the FAD of *MAOA* similarly affect MAOA function.

**Figure 2.**
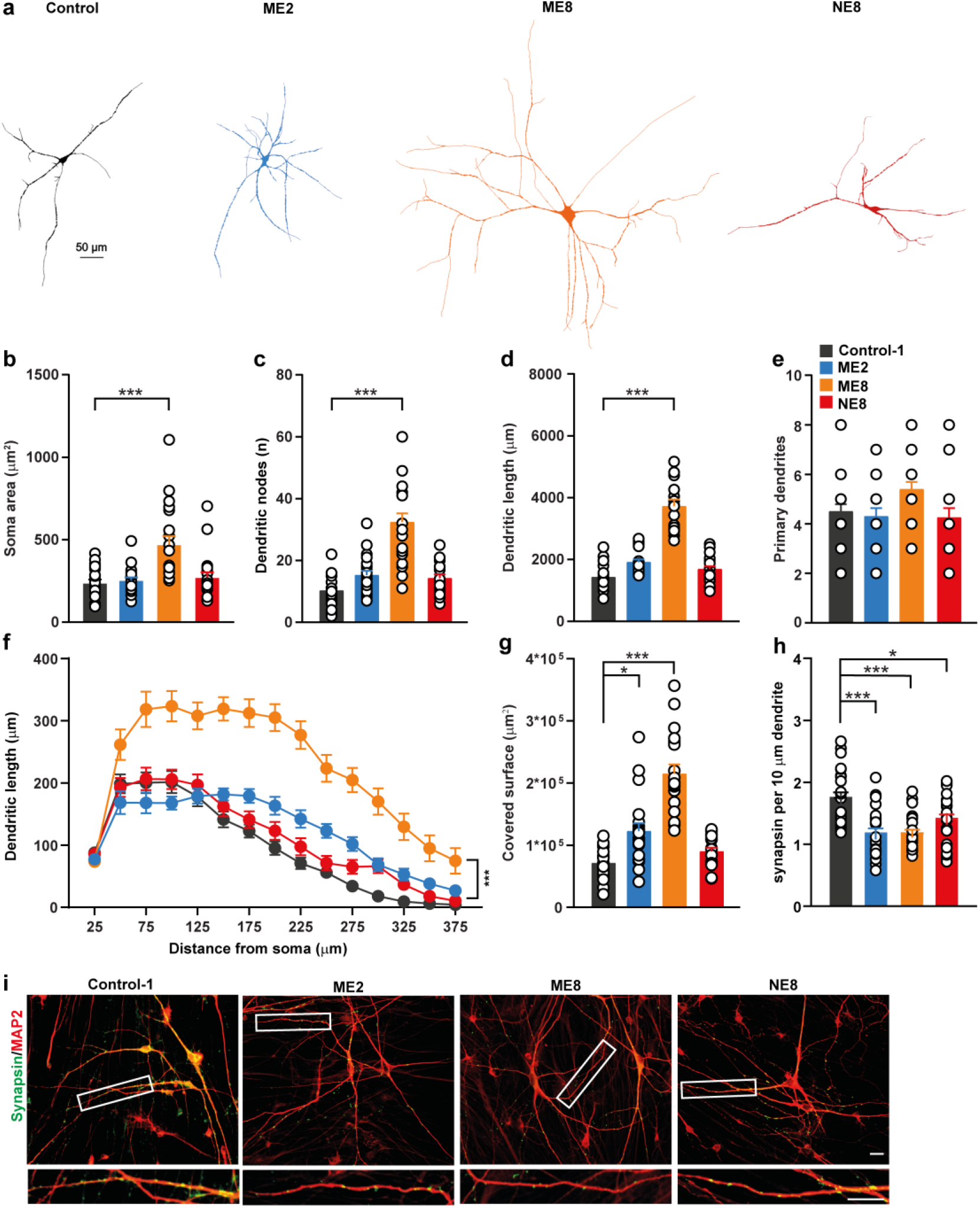
Morphological organization and synapse density of DA neurons. (a) Representative images of reconstructed DA neurons at 73 days of differentiation (DIV 73). (b-g) Parameters derived from the somatodendritic compartment. Sample size: N=20 for all lines across 3 independent differentiations). (h) Quantifications of synapse density (N positive synapsin puncta/10 μm, Sample size: Control-1 N=25, ME2 N=23, ME8 N=25, NE8 N=26. (i) Representative images of control and patient DA neurons at DIV 73 immunostained for microtubule associated protein 2 (MAP2) (red) and synapsin1/2 (green), scale bar = 20 μm. Inset shows a single stretch of dendrite (red) with synapses (green), scale bar = 10 μm. All data is represented as mean ± SEM. One-Way ANOVA with Dunnett’s correction for multiple testing was used to compare between patient lines and control lines in all parameters except Sholl analysis, where MANOVA with Bonferroni correction was used with distance and genotype as factors. *\*P<0.05; ***P<0.001*.

In addition to alterations in dendritic complexity(39, 40), neurodevelopmental disorders have been associated with synaptic deficits in rodents and humans(41). We therefore estimated synapse density using the number of presynaptic synapsin1/2 puncta per section of postsynaptic dendrite. We found that synapse density was significantly decreased in DA neurons from all three Brunner syndrome-derived lines compared to DA neurons from healthy controls at DIV73 (**Figure 2h, i**). This suggests that, whilst the effect of MAOA dysfunction on DA neuron morphology might be mutation- and/or patient specific, MAOA dysfunction induced reduction of synapse density is a general feature of DA neurons in Brunner syndrome.

### Brunner syndrome-derived DA neurons show increased neuronal network activity

Differences in network activity and organization have been observed in the brain of individuals with neurodevelopmental disorders(42), and changes in synapse density have been shown to underlie these neuronal network changes(40, 43). To investigate the neuronal network phenotypes by means of recording extracellular spontaneous activity at the population level, we generated control and Brunner syndrome DA neuron cultures grown on 6-well MEAs (**Figure 3a**). We recorded the network activity at DIV 73, the same *in vitro* timepoint at which the reduced synapse density was observed, and compared control DA neuron networks with patient networks. At this timepoint, neuronal cultures generated spontaneous activity (**Figure 3b-d**), and control lines showed similar, albeit sparse, activity levels (control-1 and control-2, **Supplementary Figure 4**). Since we detected no difference in neuronal activity between the ME2 and ME8 lines, and our data on synapse density suggest that dysfunction of the FAD domain similarly affects MAOA function in these lines (**Supplementary Figure 4**), we grouped data from the missense mutation lines for statistical analysis. Control networks mainly showed sporadic random spiking activity (**Figure 3b, e**), whereas synchronous activity at either the single electrode level (burst activity, **Figure 3f**) or throughout the entire culture (network burst activity, **Figure 3g**) was largely absent. By contrast, Brunner syndrome DA neuronal networks showed significantly higher random spiking activity at DIV 73 compared to control (**Figure 3e**). Moreover, in Brunner syndrome networks, activity occurred organized into readily observable synchronous events (network bursts) composed of many spikes occurring close in time and across the culture (**Figure 3b-c, g**). This indicates that Brunner syndrome DA neurons are more strongly integrated into a spontaneously active network than control neurons at DIV 73.

**Figure 3.**
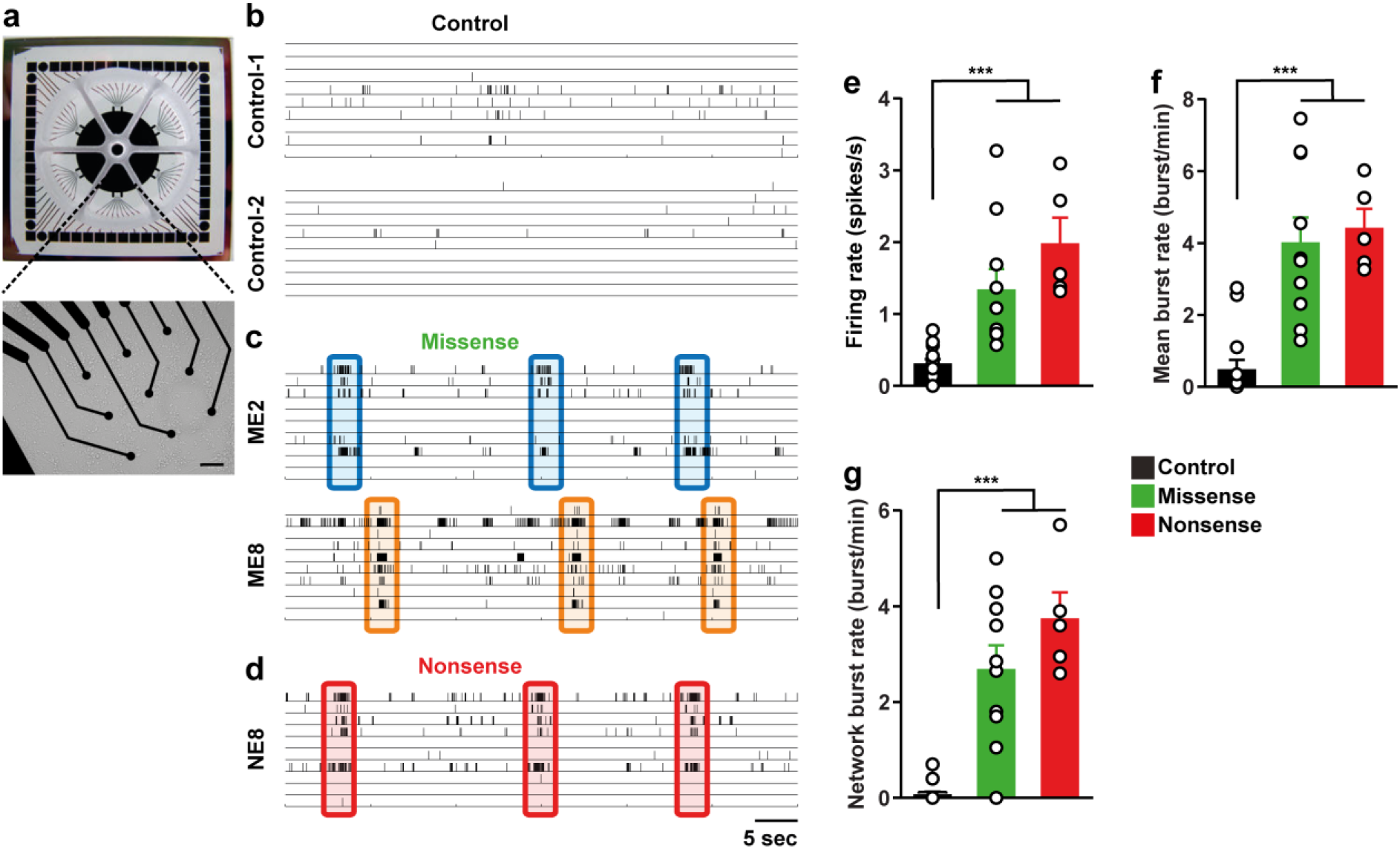
Increased neuronal network activity in Brunner syndrome DA neurons. (a) Example picture of a 6-well MEA. Each chamber is fitted with 9 recording electrodes and separated by a silicon nonconductive wall. (b-d) 60 second example raster plots of spontaneous electrophysiological activity on MEAs with DA neuron cultures at 73 days of differentiation from either healthy controls (b), or individuals with a monoamine oxidase A (*MAOA*) missense mutation (c) or nonsense mutation (d). Detected spikes are indicated as black bars. Network wide bursting activity is highlighted by colored boxes. (e) Quantification of mean firing rate. (f) Quantification of mean burst rate (g) Quantification of network burst rate. Sample size: control N=14, missense N=10, nonsense N=5. All data represent means ± SEM. One-Way ANOVA with Dunnett’s correction for multiple testing was used to compare between patient lines and control lines. *\*\*\*p<0.001*.

### MAOA dysfunction does not affect intrinsic properties and AMPAR-mediated synaptic transmission

We hypothesized that the increased network activity in Brunner syndrome DA neurons might be caused by changes in intrinsic properties of our DA neurons. We used whole-cell patch clamp to investigate passive and active intrinsic membrane properties of DA neurons, which are a measure of neuronal development and neuronal health(44). All DA neurons generated action potentials upon positive current injection to the cell soma (**Figure 4a**). At DIV 73, membrane capacitance, membrane resistance and membrane resting membrane potential were comparable across all cell lines, indicating that all assessed DA neurons showed comparable ion channel expression and level of maturity (**Figure 4b-d**). Furthermore, active (related to the action potential) properties were comparable between control and patient neurons as well (**Figure 4e-g**). These data suggest that the cell autonomous excitability and the intrinsic properties of DA neurons are not affected by mutation of *MAOA*.

**Figure 4.**
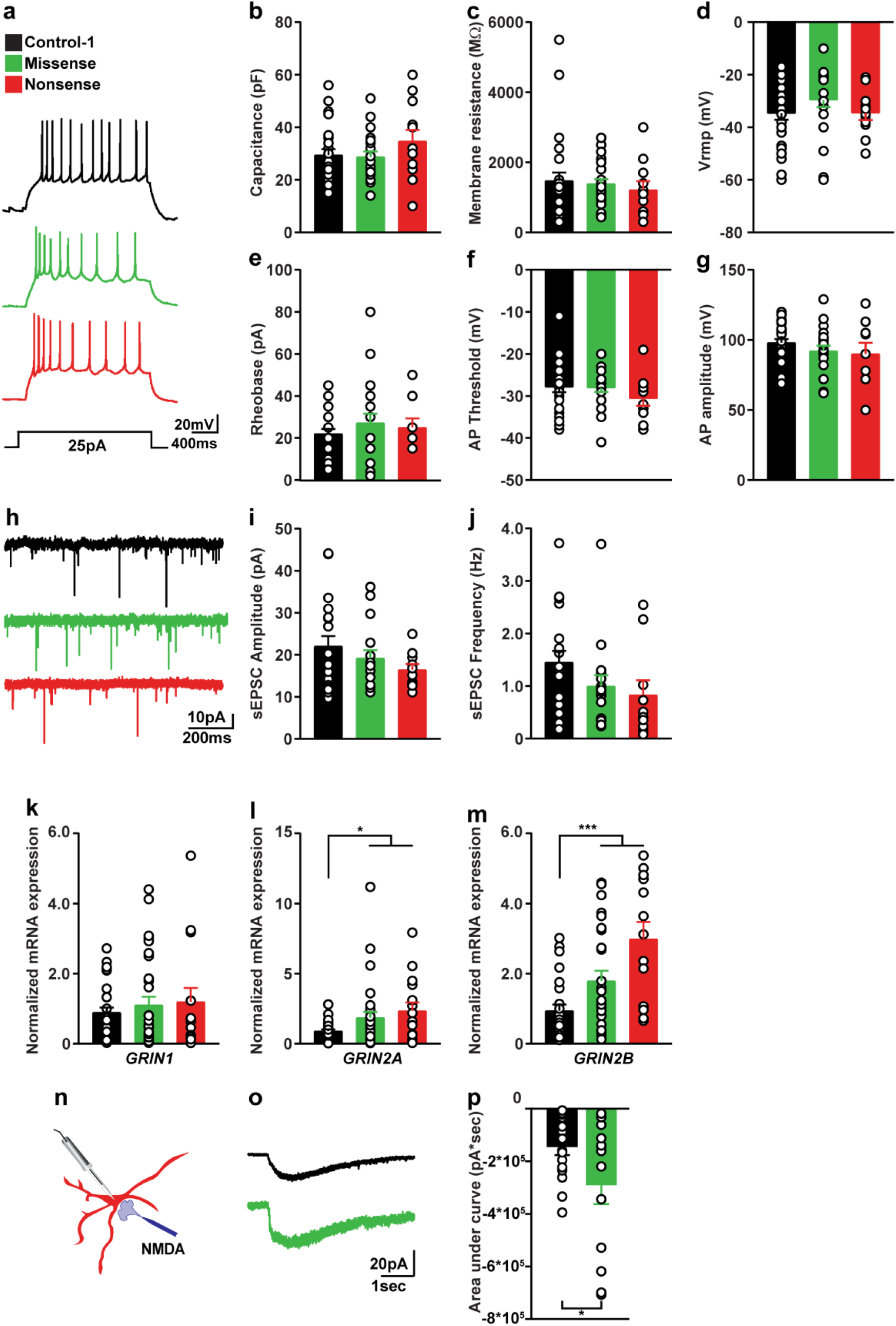
Monoamine oxidase A dysfunction results in increased N-Methyl-D-Aspartate (NMDA) mediated excitatory currents. (a) Representative traces of action potentials generated in control and patient-derived DA neurons at 73 days of differentiation. (b-d) Quantifications of passive intrinsic properties in control and patient DA neurons. (e-g) Quantifications of active intrinsic properties in control and patient DA neurons at. Sample size: control N=27, missense N=21. nonsense N=12. (h) Representative traces of spontaneous excitatory postsynaptic current (sEPSC) activity in control- and patient-derived DA neurons at DIV 73. (i,j) Quantification of spontaneous excitatory postsynaptic current (sEPSC) amplitude and frequency. Sample size: control N=19, missense N=16, nonsense N=10. (k-m) Quantification of mRNA expression of the NMDAR subunits NR1 (*GRIN1*), NR2A (*GRIN2A*) and NR2B (*GRIN2B*). Sample size: control *GRIN1* N=32, *GRIN2A* n=29, *GRIN2B* n=25. Missense *GRIN1* N=31, *GRIN2A* N=31, *GRIN2B* N=26. Nonsense *GRIN1* N=15, *GRIN2A* N=15, *GRIN2B* N=13. (n) schematic representation of NMDA receptor activation experiment. (o) Example traces of the current response to exogenous application of a high dose of NMDA (100 ms, 10 mM) at a distance of 10-20 μm from the cell soma. (p) The area under the curve (total current transfer) in control and patient lines subjected to exogenous NMDA application. Sample size: control N=16, missense N=13. All data represent means ± SEM. One-Way ANOVA with Dunnett’s correction for multiple testing was used to compare between control and patient lines. *\*P<0.05, ***P<0.001*.

(45–47). The reduced synapse density in all Brunner syndrome DA neuron lines and the increased network activity on the MEA suggest that synaptic transmission might be affected by MAOA dysfunction. Therefore, we explored whether α-amino-3-hydroxy-5-methyl-4-isoxazolepropionic acid receptor (AMPAR)-mediated spontaneous excitatory postsynaptic currents (sEPSCs) are altered in DIV 73 Brunner syndrome DA neurons. Neither sPESC frequency nor amplitude were affected between control and Brunner syndrome DA neurons (**Figure 4h-j**). Consistent with this, we did not find a significant change in mRNA expression of the most common AMPAR subunits (*GRIA1-4*) across lines (**Supplementary Figure 5**). We did observe an increase in mRNA expression of the *GRIA1* subunit in the NE8 line, but this was not reflected by a change in AMPAR-mediated currents. Taken together, these data show that AMPAR-mediated currents are not affected by MAOA dysfunction.

### MAOA dysfunction leads to NMDAR hyperfunction

Next to AMPAR mediated currents, NMDAR-mediated currents are an important component of balanced network activity both *in vitro* and *in vivo*, and changes in NMDAR function have been shown to affect network function in hiPSC-derived neuronal cultures(33, 48). We therefore hypothesized that aberrant NMDAR function could be responsible for the hyperactive network phenotypes in the Brunner syndrome DA neurons. To test this, we measured the transcripts of the most common NMDAR subunits by RT-qPCR for all hiPSC derived DA neuron lines. We found no significant changes in *GRIN1* mRNA expression, which codes for the mandatory subunit present in functional NMDARs (**Figure 4k**). However, we found a two-fold upregulation of *GRIN2A* and *GRIN2B* mRNA, which encode NMDAR subunit 2A and subunit 2B, respectively (**Figure 4l-m**). Aberrant expression of *GRIN2A* and *GRIN2B* has been shown to directly affect NMDA mediated current responses(49, 50). To test whether the increased NMDAR subunit expression leads to increased NMDAR-mediated currents, control and missense Brunner syndrome DA neurons were stimulated with a local exogenous application of NMDA We found that the total current transfer mediated by the NMDA application was significantly increased in these neurons compared to controls (**Figure 4o-p**). Taken together, this suggests that increased NMDAR expression or function might underlie the increased network activity observed in Brunner syndrome DA neuronal networks at DIV 73.

### Correction of *MAOA* mutation restores NMDAR expression and DA neuronal network activity

Our data suggest that the increase in *GRIN2A* and *GRIN2B* expression in the patient lines, and the concomitant increase in NMDA mediated currents in the ME2 and ME8 missense lines, are a direct consequence of *MAOA* mutation. In order to further validate this causality, we generated an isogenic hiPSC line (ME8-CRISPR) in which we corrected the p.C266F mutation present in the ME8 line through CRISPR/Cas9 mediated homologous recombination (**Figure 5a, Supplementary Figure 6**). We found that NMDAR subunit transcript levels in ME8-CRISPR DA neurons were similar to control values at DIV 73 (**Figure 5b-d**). This further indicates that normal MAOA activity is essential for the regulation of *GRIN2A* and *GRIN2B* expression. Correction of the ME8 missense mutation also resulted in the normalization of the NMDA-induced NMDAR-mediated current transfer (**Figure 5e, f**). Finally, we found that restoration of MAOA function also normalized activity on the MEA to control values (**Figure 5g**); whereas ME8 DA neurons showed increased network activity, this increase was absent in ME8-CRISPR DA neuronal networks, as the mean firing rate (**Figure 5h**) and mean burst rate (**Figure 5i**) were similar to control values. Therefore, we conclude that rescue of MAOA mutation results in normalization of *GRIN2A and GRIN2B* expression, which is reflected by a restoration of neuronal network activity to control values. This implicates aberrant expression of NMDARs in the neuronal network phenotype observed in DA neurons derived from individuals with Brunner syndrome.

**Figure 5.**
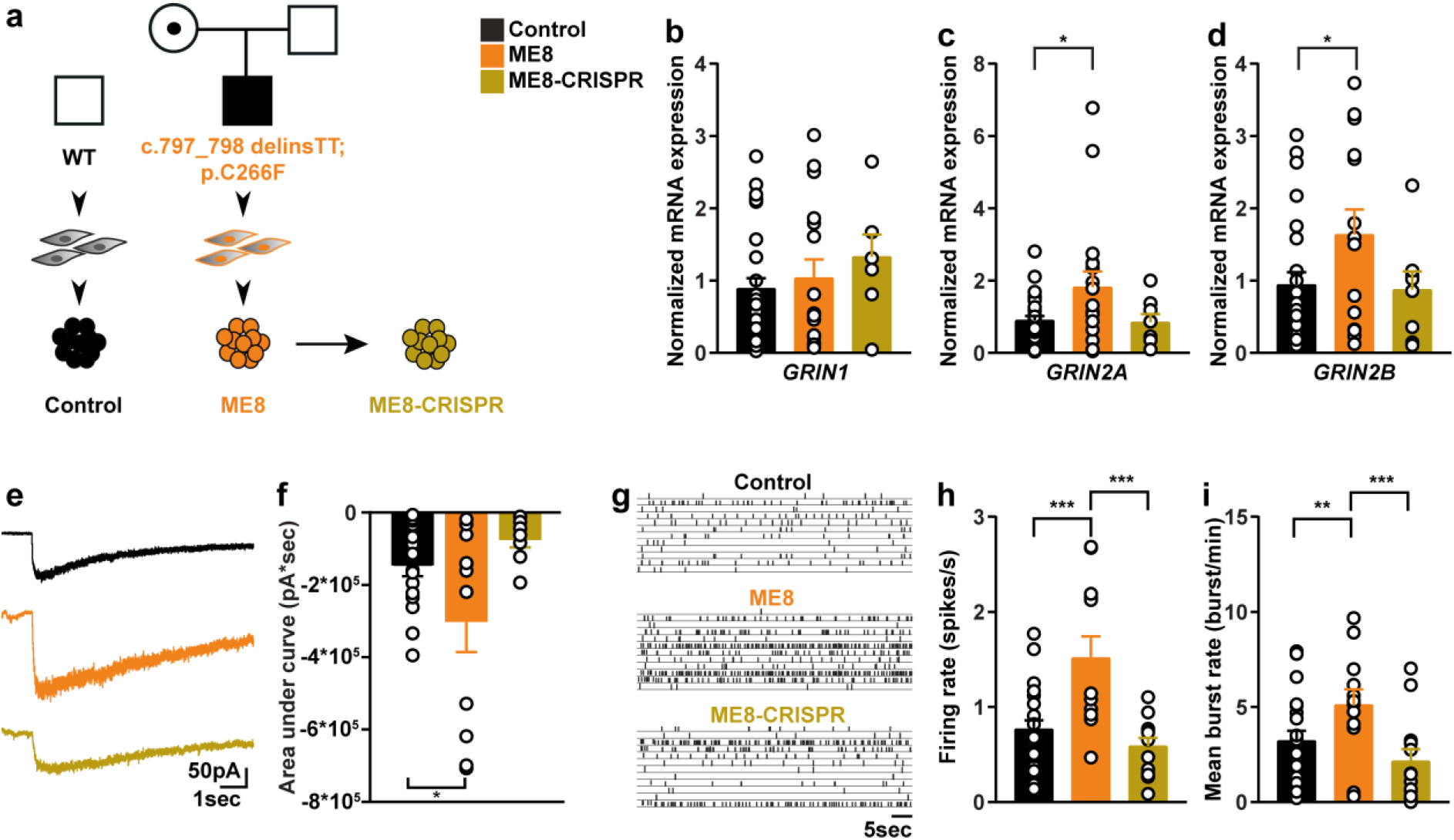
Correction of a missense mutation restores *GRIN2A* and *GRIN2B* expression, N-Methyl-D-Aspartate-mediated currents and neuronal network activity. (a) Overview of the lines used for the CRISPR/Cas9-mediated rescue of Monoamine oxidase A function. (b-d) Quantification of mRNA expression of NMDAR subunits *GRIN1, GRIN2A* and *GRIN2B*. Sample size: control *GRIN1* N=32, *GRIN2A* N=29, *GRIN2B* N=25. ME8 *GRIN1* N=16, *GRIN2A* N=18, *GRIN2B* N=12. ME8-CRISPR *GRIN1* N=8, *GRIN2A* N=7, *GRIN2B* N=8 from at least three different neuronal preparations. (e) Example traces of the current response to exogenous NMDA application in control, ME8 and ME8-CRISPR DA neurons at 73 days of differentiation (DIV73). (f) Total current transfer (area under the curve) upon NMDA application (*P=0.0219* between control and ME8) at DIV 73. Sample size: control N=16, ME8 N=12, ME8-CRISPR N=13. (g) 60 second example trace of spontaneous network activity recorded on a 24-well microelectrode array system in control, ME8 and ME8-CRISPR DA neuronal cultures at DIV 69. (h) Quantification of mean firing rate (*P=0.00044* between control and ME8, and *P=0.00011* between ME8 and ME8-CRISPR). (i) Quantification of mean burst rate (*P=0.0048* between control and ME8 and *P=0.0005* between ME8 and ME8-CRISPR) at DIV 69. Sample size: control N=23, patient ME-8 N=12, ME8-CRISPR N=13. One-Way ANOVA with Dunnett’s correction for multiple testing was used to compare control, ME8 and ME8-CRISPR. Data are shown as mean ± SEM.\**P<0.05,**P<0.01,***P<0.001*.

## Discussion

Monoamine aberration through dysfunction of MAOA results in Brunner syndrome. Although the syndrome has been described almost three decades ago, insight into the molecular mechanisms of the disorder is still lacking. Here, we developed a hiPSC-derived DA neuron model to assess the molecular and cellular phenotypes underlying brain dysfunction in Brunner syndrome. Until now, only four families have been reported in which individuals have Brunner syndrome(5–8). We were able to include individuals from three of these families into our study. hiPSC-derived DA neurons generated from individuals with Brunner syndrome showed reduced synapse density but increased network activity. The phenotype could be linked to increased *GRIN2A* and *GRIN2B* expression and NMDAR hyperfunction. Lastly, we were able to restore DA neuronal network activity and NMDAR-mediated activity to control values by correcting a missense mutation using CRISPR/Cas9.

One of the opportunities of hiPSC technology is that patient-specific mutations can be investigated using patient-derived cell lines. In our case, we found that lines from the different individuals with Brunner syndrome exhibit both overlapping and distinct phenotypes. As such, some differences between control and patient-derived DA neurons could not be reliably attributed to MAOA dysfunction. For example, DA neuron dendritic morphology between lines derived from individuals with and without Brunner syndrome was highly cell line specific, and we suspect that the differences we found between control and Brunner syndrome lines were strongly affected by the individual genetic background rather than being a clear consequence of MAOA dysfunction. This contrasts with conclusions drawn from previous studies of hemizygous *Maoa* knockout mice, in which increased dendritic arborization of pyramidal neurons in the orbitofrontal cortex was described(51). However, it is not known whether affected neurons in mouse orbitofrontal cortex expressed MAOA. As such, the changes in neuronal morphology in these mice do not necessarily reflect an effect of cell autonomous reduction of MAOA expression. Instead, it is possible that differences in monoamine levels in these mice during brain development can affect neuronal morphology, as monoamines such as dopamine and serotonin can induce neurite growth in rodent neurons(13, 52, 53). Additionally, our human DA neuron cultures are dependent on glial support from wildtype rodent astrocytes, which express both MAOA and MAOB(54) and are highly involved in the maintenance of dopamine levels *in vivo*(38). As such, the influence of MAOA dysfunction in the human DA neurons on extracellular dopamine levels in our *in vitro* cultures might be limited, which could lead to occlusion of somatodendritic phenotypes. Recent developments that enable the generation of homogenous cultures of hiPSC derived astrocytes(55) offer exciting opportunities to further explore the contribution of non-neuronal MAOA dysfunction to monoamine aberration *in vitro*.

Recent single cell RNAseq profiling confirms that *MAOA* is not exclusively expressed in DA neurons, but is expressed in neural progenitor cells and other monoaminergic neurons(26, 56). Similar to aberrant dopamine signaling, dysfunction of serotonergic systems is associated with aggression and impulsivity(23, 57). We constrained our investigation to MAOA dysfunction in a homogenous DA neuron population, where MAOA dysfunction shows a clear neuronal phenotype. It will be interesting to extend these investigations to include other monoaminergic neuron types, and established protocols to generate a homogenous population of hiPSC-derived serotonergic neurons(37, 58). Thus, it is possible to assess whether the same molecular mechanisms affected by MAOA dysfunction in DA neurons can be generalized to other neuron subtypes in which MAOA is expressed.

The individuals with Brunner syndrome all carry rare mutations, which lead to either complete loss or reduced activity of the MAOA enzyme (5–7, 35). In the general population, the VNTR polymorphism in the promoter of *MAOA* can induce different levels of transcriptional activity(36). Low activity alleles of *MAOA* (*MAOA-L*) has been associated with increased anti-social behavior in individuals subjected to childhood maltreatment(59, 60). Interestingly, we recently showed increased structural and functional connectivity of brain regions associated with emotion regulation in healthy individuals carrying *MAOA-L* alleles compared to those carrying high activity *MAOA* (*MAOA-H*) alleles using magnetic resonance imaging(61). The individuals with Brunner syndrome included here carry both *MAOA-L* (ME8) and *MAOA-H* (ME2 and NE8). Intriguingly, the ME8 DA neurons did show an increase in dendritic complexity compared to both controls and the ME2 and NE8 patient lines. However, this did not result in differences in the functional network phenotype between ME8 and the other lines. Furthermore, ME2, ME8 and NE8 all showed similar synapse densities and *GRIN2A* and *GRIN2B* expression. This suggests that not all functional phenotypes shown by MAOA dysfunction are affected by the presence of *MAOA-L* or *MAOA-H* VNTRs. The increased dendritic complexity might be a consequence of MAOA dysfunction aggravated by the presence of the *MAOA-L* VNTR. Further exploration of how low and high activity *MAOA* alleles affect DA neuron function in healthy subjects can help us understand molecular mechanisms regulated by MAOA.

Similar to the effect of MAOA dysfunction in human DA neurons, increased expression of the *GRIN2A* and *GRIN2B* NMDAR subunits has been observed in prefrontal cortex of *Maoa* hemizygous knockout mice. The prefrontal cortex is a highly heterogenous region, and until now it was unclear whether the changes in NMDAR expression and NMDA mediated currents that were observed in *Maoa* hemizygous knockout mouse were established through cell-autonomous mechanisms. Isogenic rescue of MAOA mutation in the ME8 line results in normalization of *GRIN2A* and *GRIN2B* expression and NMDAR-mediated currents to control levels, which suggests that MAOA dysfunction cell-autonomously affects NMDAR activity. Importantly, the restoration of NMDAR function results in reversal of the network phenotype in DA neuron cultures, which could also be an explanation for the positive effects of NMDAR antagonism on locomotor behavior of *Maoa* hemizygous knockout mice(19). The overlap in mechanisms affected in the prefrontal cortex of these mice and hiPSC-derived DA neurons of individuals with Brunner syndrome shows that DA neuron cultures are a viable *in vitro* system to investigate possible therapeutic strategies.

The protocol we describe here results in highly homogenous and reproducible cultures of DA neurons. Growing hiPSC-derived DA neuron networks on MEAs enables us to study patient-specific neuronal networks. Most investigations that use hiPSC-derived DA neurons have focused on cell-based therapy for neurodegenerative disorders such as Parkinson’s disease, lacking characterization of network activity(62, 63). In one study, hiPSC derived DA neurons have been cultured on MEAs to investigate neuronal phenotypes in a monozygotic twin pair discordant for Parkinson’s disease(64). Reduced network activity was seen in DA neurons derived from the twin with Parkinson’s disease, which shows that network phenotypes of hiPSC derived DA neurons can be characterized using MEAs. In our study, we combine an extensive characterization of both the single-cell and network properties of hiPSC-derived DA neurons and how these can be related to differences at the molecular level in neurodevelopmental disorders with a monogenic cause.

In conclusion, our data suggest that MAOA dysfunction affects DA neuron function in individuals with Brunner syndrome and that NMDAR hyperfunction is a key contributor to network dysfunction in Brunner syndrome DA neuron cultures. These alterations on the network level might be able to explain parts of Brunner syndrome associated impulsivity and maladaptive externalizing behavior. Manipulation of NMDAR function could be a viable opportunity toward the development of possible therapeutic strategies.

## Acknowledgements

The authors would like to acknowledge support from the Netherlands Organization for Scientific Research (NWO) Vici Innovation Program (grant 016-130-669 to BF), from the European Community’s Seventh Framework Programme (FP7/2007–2013) under grant agreement no. 602805 (Aggressotype), and from the European Community’s Horizon 2020 Programme (H2020/2014–2020) under grant agreements no. 667302 (CoCA) and no. 728018 (Eat2beNICE). Additional support is received from the Dutch National Science Agenda for the NeurolabNL project (grant 40017602) the NWO grant 012.200.001 and 91217055 (to N.N.K.), SFARI grant 610264 (to N.N.K), ERA-NET NEURON DECODE! grant (NWO) 013.18.001 (to N.N.K) and ERA-NET NEURON-102 SYNSCHIZ grant (NWO) 013-17-003.4538 (to D.S).

## Author contributions

Y.S., J.v.R., B.F. and N.N.K. conceived and supervised the study. Y.S., J.v.R., B.F. and N.N.K. designed all the experiments. Y.S. performed all cell culture, generated the isogenic CRISPR/Cas9 mediated lines and acquired all MEA data. J.v.R. performed all single-cell electrophysiological experiments, M.B. performed neuronal reconstruction. B.M., M.H., T.K.G., C.S. performed additional experiments. M.F., S.K-S, D.S., H.B., L.F. and E.P. provided resources. Y.S., J.v.R., M.B., B.M. performed data analysis. Y.S., J.v.R., B.F., and N.N.K. wrote the paper. D.S., H.B., and B.F. edited the paper.

