## Supplementary methods, tables, and figures for "Brunner syndrome associated MAOA dysfunction in human dopaminergic neurons results in NMDAR hyperfunction and increased network activity"

Running title: **MAOA deficiency leads to increased neuronal activity**

Yan Shi[1*], Jon-Ruben van Rhijn[2*], Maren Bormann[2], Britt Mossink[1,2], Monica Frega[1,3], Marina Hakobjan[1], Teun Klein Gunnewiek[1,4], Chantal Schoenmaker[1], Elizabeth Palmer[5,6], Laurence Faivre[7,8], Sarah Kittel-Schneider[9,10], Dirk Schubert[2], Han Brunner[1,12], Barbara Franke[1,11#], Nael Nadif Kasri[1,2#]

*These authors contributed equally

[#]Shared final responsibility

[1]Department of Human Genetics, Donders Institute for Brain, Cognition and Behavior, Radboud University Medical Center, Nijmegen, The Netherlands
[2]Department of Cognitive Neuroscience, Donders Institute for Brain, Cognition and Behavior, Radboud University Medical Center, Nijmegen, The Netherlands
[3]Department of Clinical neurophysiology, University of Twente, 7522, NB Enschede, Netherlands
[4]Department of Anatomy, Donders Institute for Brain, Cognition and Behavior, Radboud University Medical Center, Nijmegen, The Netherlands
[5]Genetics of Learning Disability Service, Hunter Genetics, Waratah, NSW, Australia
[6]School of Women's and Children's Health, University of New South Wales, Randwick, NSW, Australia
[7]Centre de Référence Anomalies du développement et Syndromes malformatifs and FHU TRANSLAD, Hôpital d'Enfants, Dijon, France
[8]INSERM UMR1231 GAD, Faculté de Médecine, Université de Bourgogne, Dijon, France.
[9]Department of Psychiatry, Psychosomatic Medicine and Psychotherapy, University Hospital, Goethe-University, Frankfurt, Germany
[10]Department of Psychiatry, Psychosomatic Medicine and Psychotherapy, University Hospital Würzburg, Würzburg, Germany
[11]Department of Psychiatry, Donders Institute for Brain, Cognition and Behavior, Radboud University Medical Center, Nijmegen, The Netherlands
[12] Department of Clinical Genetics, MUMC+, GROW School of Oncology and Developmental Biology, and MHeNS School of Neuroscience and Maastricht University, Maastricht, the Netherlands

Corresponding author: Nael Nadif Kasri; Radboud University Medical Centre, Department of Human Genetics, Geert Grooteplein 10, 6525GA, Nijmegen, The Netherlands. Tel: +31 24 3614242,

### Supplementary methods and materials

- Cell Culture of hiPSCs
- Differentiation of hiPSCs into dopaminergic neurons
- Gene expression analysis
- Neuronal reconstruction and quantitative morphometrical analysis.
- Immunocytochemistry
- Microelectrode array and data analysis
- Single-cell electrophysiology
- Genotyping of the MAOA promoter VNTR polymorphism

### Supplementary Figures

- **Supplementary Figure 1**: Validation of human induced pluripotent stem cell generation using pluripotency markers
- **Supplementary Figure 2:** Variable number of tandem repeat (VNTR) polymorphism in the monoamine oxidase A (*MAOA*) promoter
- **Supplementary Figure 3:** Microtubule Associated Protein 2 (MAP2) staining of dopaminergic neurons at day 73 of differentiation (DIV 73)
- **Supplementary Figure 4:** Activity of dopaminergic neurons on microelectrode arrays (MEA) at day 73 of differentiation (DIV 73) separated for all control and patient lines
- **Supplementary Figure 5:** Expression of α-amino-3-hydroxy-5-methyl-4-isoxazolepropionic acid receptor subunits in control and patient lines
- **Supplementary Figure 6:** CRISPR/Cas9 mediated correction of the p.C266F (ME8) monoamine oxidase A (*MAOA*) mutation

### Supplementary Tables

- **Supplementary Table 1:** information about the cell lines used in the study, supporting Figure 1
- **Supplementary Table 2:** Primers for qRT-PCR, supporting Figure 1, 4, 5 and Supplementary Figure 4
- **Supplementary Table 3:** statistics related to Figure 1 and Supplementary Figure 2
- **Supplementary Table 4:** statistics related to Figure 2
- **Supplementary Table 5:** statistics related to Figure 3 and Supplementary Figure 4
- **Supplementary Table 6:** statistics related to Figure 4
- **Supplementary Table 7:** statistics related to Figure 5 and Supplementary Figure 6

### Supplementary methods and materials

#### Cell Culture of hiPSCs

The hiPSC line control-1 was derived from dermal fibroblasts of a male healthy volunteer(1). The iPSC line control-2 was obtained from(2) and was generated from dermal fibroblasts of a healthy male donor by episomal reprogramming(3). The iPSC lines ME2, ME8 and NE8 were derived from dermal fibroblast biopsies of male individuals diagnosed with Brunner syndrome described previously(4-7). hiPSCs were generated from fibroblasts by introduction of the Yamanaka factors *Sox2*, *Klf4*, *Oct4* and *c-Myc* via lentiviral transduction. All hiPSC reprogramming and characterization of the pluripotency markers was done by the Radboudumc Stem Cell Technology Center (SCTC) (**Supplementary Figure 1**). The hiPSCs were maintained in Essential 8 (E8) or Essential 8 flex complete medium with 100 µg/ml penicillin/streptomycin or 100 µg/ml Primocin (InvivoGen) on vitronectin coated cell culture plates (Corning) at 37˚C/5% $CO_2$. Cell colonies were passaged by 0.5 mM ultrapure EDTA treatment for 2 or 3 minutes in Dulbecco’s phosphate-buffered saline (DPBS) without calcium and magnesium. Unless indicated otherwise, all the reagents were bought from Thermo Fisher Scientific Inc.

#### Differentiation of hiPSCs into dopaminergic (DA) neurons

We split hiPSC colonies into single cells by Accutase treatment (5 min/ 37ºC) two days before the start of differentiation. 1-2*$10^4$/ $cm^2$ cells were replated on a vitronectin-coated plate in E8 complete medium with 2 µM thiazovivin (Sigma) or 1x RevitaCell Supplement. For differentiation, the E8 complete medium was replaced with Knockout DMEM from DIV 0 to DIV 5, supplemented with 15% Knockout Serum Replacement, 1x GlutaMAX, 100 U/mL Penicillin-Streptomycin and 1x MEM Non-Essential Amino Acids Solution. 0.1 mM β-mercaptoethanol was added freshly to the Knockout DMEM before each medium change. From DIV 6 to DIV 9, the Knockout DMEM medium was gradually replaced by 25%, 50%, 75%, and 100% N2 medium (DMEM/F12 with 1x N-2 supplement). DA neurons were cultured in N2 medium until DIV 11. From DIV 12 on, the DA neurons were cultured in neurobasal medium supplemented with 1x B27 and 1x GlutaMAX and half medium was refreshed every two days. During the differentiation and lineage specification, different combinations of small molecules and/or growth factors were added at different time points: 10 µM LDN-193189 (Stemgent Inc.) from DIV 1 to DIV 11; 10 µM SB431542 (Stemgent Inc.) from DIV 1 to DIV 5; 2 µM Purmorphamine (Stemgent Inc.) and 100 ng/mL recombinant human fibroblast growth factor 8a (FGF-8a, R&D system) from DIV 2 to DIV 11; 3 µM CHIR99021 (Stemgent Inc.) from DIV 3 to DIV 12; 100 ng/ml recombinant human sonic hedgehog (C24II, SHH, R&D system) from DIV 12 to the end of the differentiation. 20 ng/ml recombinant brain-derived neurotrophic factor (BDNF, Peprotech), 20 ng/ml recombinant glial-derived neurotrophic factor (GDNF, Peprotech), 0.5 mM adenosine 3',5'-cyclic monophosphate (cAMP, Enzo Life Science), 2 ng/ml transforming growth factor beta 3 (TGFβ3, Millipore), 200 µM ascorbid acid (AA, Sigma) and 10 nM γ-secretase inhibitor IX (DAPT, Millipore, Calbiochem) were added into the medium from DIV 11 to the end of differentiation. The cells were passaged only when they were 100% confluent using accutase treatment. At DIV 20, DA neurons were split into single cells and 1-2x $10^4$/$cm^2$ cells were replated on a poly-L-ornithine (Sigma, 50 µg/ml) and murine Laminin (Sigma, 10 µg/ml) double-coated plate. From DIV 22 on, 10 µM DAPT was used to promote DA neuron maturation. Rat astrocytes [prepared as previously described(8)] were cocultured with DA neuron progenitors from DIV 24 to promote maturation.

**Gene expression analysis**

RNA was isolated from hiPSCs and differentiated DA neurons with the RNeasy Mini Kit (Qiagen) according to the manufacturer’s instructions. 0,5-1 µg RNA was retro-transcribed into cDNA by the iScript cDNA Synthesis Kit (Bio-Rad Laboratories, Inc) according to the manufacturer’s instructions. The gene expression profile of hiPSCs and differentiated DA neurons was measured by quantitative real-time PCR (qRT-PCR) using the Applied Biosystems 7500 Fast RT-PCR System. Used qRT-PCR primers are listed in **Supplementary Table S1**. Beta-2-Microglobulin (B2M) was used as reference gene. The Ct value of each target gene was normalized against the Ct value of the reference gene [ΔCt= [Ct(target)-Ct(B2M)]. The relative expression was calculated as $2^{-\Delta\Delta Ct}$ and represented as fold change of gene expression when compared to corresponding control conditions [$2^{-\Delta\Delta Ct}=2^{\Delta Ct(target)-\Delta Ct(control)}$].

**Neuronal reconstruction and quantitative morphometrical analysis.**

Widefield fluorescent images of MAP2-labelled hiPSC-derived dopaminergic neurons were taken at 20x magnification using a Zeiss Axio Imager Z1 with apotome (Carl Zeiss AG, Germany). The images were stitched using Fiji software and the somatodendritic domains of individual neurons were reconstructed using Neurolucida 360 (Version 2017.01.4, Microbrightfield Bioscience, Williston, USA). Only neurons with at least two primary dendrites and at least one dendritic branch point were selected for reconstruction and further analyses. Sholl analysis(9) was used to determine the dendritic length of the neurons within a series of concentric circles at 25 µm intervals from the soma. All the morphological data were acquired and analyzed blind to the genotype of the neurons. For the somatodendritic properties and Sholl analysis, the significance was determined by using Wilks' Lambda multivariate analysis of variance (MANOVA) followed by the Bonferroni post-hoc (IBM SPSS Statistics, version 24.0, IBM, Armonk, USA).

**Immunocytochemistry**

Cells plated on coverslips were fixed with 4% paraformaldehyde/4% sucrose (v/v) (Sigma) in PBS for 15 min at room temperature (RT). Non-specific binding was avoided by incubation in 5% normal goat serum (Thermo Fisher Scientific)/ 0.4% Triton X-100(Sigma)/1% Glycine (Sigma) in PBS (blocking solution) at RT for one hour. The primary and secondary antibodies were diluted in the blocking solution and applied overnight at 4°C or 1h at RT respectively.

Cell nuclei were stained with Hoechst 33342 (Molecular probes), and the coverslips were mounted with DAKO fluoromount medium (Agilent). The primary antibodies used were: mouse anti-MAP2 (1:1000; Sigma M4403), guinea pig anti-MAP2 (1:1000; Synaptic Systems 188004), guinea pig anti-synapsin 1/2 (1:1000; Synaptic Systems 106004), or mouse anti-TH (1:200; Sigma TH-16). Secondary antibodies used were: goat anti-guinea pig Alexa Fluor 568 (1:1000, Invitrogen A-11075), goat anti-rabbit Alexa Fluor 488 (1:1000, Invitrogen A-11034), goat anti-mouse Alexa Fluor 488 (1:1000, Invitrogen A-11029), or goat anti-mouse Alexa Fluor 568 (1:1000, Invitrogen A-11031). DA neurons were imaged at 63x magnification using a Zeiss Axio Imager Z1 with apotome. Synapse density was assessed through manual counting using Fiji software.

**Microelectrode array(MEA) and data analysis**

Neuronal network activity was measured using 6-Well or 24-well microelectrode array devices (Multichannel Systems, MCS GmbH, Reutlingen, Germany). DA progenitor cells (DIV 20) were plated on MEAs and further cultured as previously described. Each well was comprised of either 9 (6-Well MEA), or 12 recording electrodes (24-well MEA) and a grounding electrode. Activity in 6-well MEAs was recorded for 20 min using the MEA60 System (MCS GmbH, Reutlingen, Germany) as described before(8) with a high pass filter (Butterworth, 100 Hz cutoff frequency). Recordings on the 24-well MEA system were conducted as described before(10). During all recordings for the 6-well and 24-well MEAs, the temperature was maintained at 37°C and with continuous flow of humidified carbogen (95% $O_2$, 5% $CO_2$). Data was sampled at 10 kHz through either MC-Rack software (6-well MEA) or Multiwell-Screen software (24-well MEA) (MCS GmbH, Reutlingen, Germany).

Data analysis from 24-well MEAs was performed off-line by using Multiwell Analyzer (MCS GmbH, Reutlingen, Germany, i.e. software from the 24-well MEA system that allows the extraction of the filtered output data per electrode) and in-house algorithms to average all the data per well as previously described (8, 11). The extracted parameters encompassed the mean firing rate (MFR, spikes/second) and mean burst rate (MBR, bursts/min). The MFR was computed by averaging the firing rate of each channel, which is averaged for all the active electrodes of the MEA. Spikes were grouped into a burst if at least 4 consecutive spikes were detected with a smaller than 30 milliseconds inter-spike-interval. All bursts were merged that were less than 65 milliseconds apart. Bursts with a duration of lower than 50 milliseconds were removed from analysis.

For 6-well MEA data analysis, spikes and bursts were detected by using the Precise Timing Spike Detection algorithm (PTSD)(12) and the Burst Detection algorithm(13) embedded in the SpyCode software(11). The mean firing rate (spikes/s) of the network was computed by averaging the firing rate of each channel, which is averaged for all the active electrodes (MFR>0.1Hz) of the MEA. The burst was computed as at least 5 spikes in a burst with a maximum inter-spike-interval of 80 milliseconds. The network burst was defined as synchronized bursts that occurs in >50% of the active channels. The network burst rate (burst/min) was calculated as the amount of network bursts per minute. Statistical analysis was conducted in PRISM (Graphpad PRISM 7.0, Graphpad Software, San Diego, CA).

**Single-cell electrophysiology**

All single-cell electrophysiological recordings were conducted on DIV 73 DA neurons. Coverslips plated with neurons were transferred to a recording chamber continuously perfused with oxygenated (95% $O_2$ / 5% $CO_2$) and heated (32 °C) recording artificial cerebrospinal fluid (ACSF) containing (in mM): 124 NaCl, 3 KCl, 1.25 $NaH_2PO_4$, 2 $CaCl_2$, 1 $MgCl_2$, 26 $NaHCO_3$, 10 Glucose. Patch pipettes with filament (5.5-7.5 MΩ) were made from borosilicate glass capillaries (Science Products GmbH, Hofheim, Germany). For all recordings of intrinsic properties and spontaneous activity, a potassium-based intracellular solution containing (in mM) 130 K-Gluconate, 5 KCl, 10 HEPES, 2.5 $MgCl_2$, 4 $Na_2$-ATP, 0.4 $Na_3$-ATP, 10 Na-phosphocreatine, 0.6 EGTA (pH 7.2 and 290 mOsmol) was used. The resting membrane potential (Vrmp) was measured immediately after generation of a whole cell configuration. All other measurements were conducted at a holding potential of -60 mV). Passive membrane properties were determined via voltage step of -10 mV. Active intrinsic properties were measured with a stepwise current injection protocol.

For recording of current changes upon exogenous NMDA application coverslip were placed in $MgCl_2$-free ACSF, a cesium-based intracellular solution containing (in mM) 115 $CsMeSO_3$, 10 CsCl, 10 HEPES, 2.5 $MgCl_2$, 4 $Na_2ATP$, 0.4 NaGTP, 10 Na-Phosphocreatine; 0.6 EGTA, 10 QX-314 was used. Application pipettes (2-4 MΩ) were made from borosilicate glass with filament. NMDA (10 mM) was dissolved in ACSF and locally applied using a pressure ejection system (PDES-2DZ, NPI, Tamm, Germany). The ejection pressure was set to 5 psi/0.4 bar and injection duration was set to 100 ms. CNQX (5µM, Tocris, Bristol, United Kingdom) was used to block all AMPAR-mediated currents. Cells were visualized with an Olympus BX51WI upright microscope (Olympus Life Science, PA, USA), equipped with a DAGE-MTI IR-1000E (DAGE-MTI, IN, USA) camera). Activity was recorded using a Digidata 1440A digitizer and a Multiclamp 700B amplifier (Molecular Devices). Sampling rate was set at 20 kHz (voltage measurements) or 10 kHz (current measurements) and a lowpass 1 kHz filter was used during recording. Recordings were not corrected for liquid junction potential (± 10 mV). Series resistance was monitored on-line and cells were discarded if series resistance increased above 1:10 of membrane resistance. Intrinsic electrophysiological properties were analysed using Clampfit 10.7 (molecular devices, CA, USA), and sPSCs were analysed using MiniAnalysis 6.0.2 (Synaptosoft Inc, GA, USA).

**Genotyping of the *MAOA* promoter VNTR polymorphism**

The genomic DNA was amplified with 1x AmpliTaq Gold® 360 Master Mix (Life Technologies) and 0.33 mM of fluorescently labeled forward primer (FAM-5’-

ACAGCCTGACCGTGGAGAAG-3′) and reverse primer (5′-GAACGGACGCTCCATTCGGA-3′) in a total volume of 7.5 µl using the protocol: 95 $^{o}$C for 10 min followed by 35 cycles of denaturation for 30 s at 95 $^{o}$C, 30 s annealing at 60 $^{o}$C, primer extension at 72 $^{o}$C for 1 min, and a final extension at 72 $^{o}$C for 10 min. Fragment length analysis of the PCR product was performed by an automated capillary sequencer ABI3730 (Applied Biosystems, Nieuwerkerk a/d Ijssel, The Netherlands) using standard conditions (1 µl of the 1:20 diluted PCR product together with 9.7 µl formamide and 0.3 µl GeneScan-600 Liz Size Standard TM (Applied Biosystems). Results were analyzed with GeneMarker version 2.6.7 (SoftGenetics, US).

**Supplementary Figure 1**

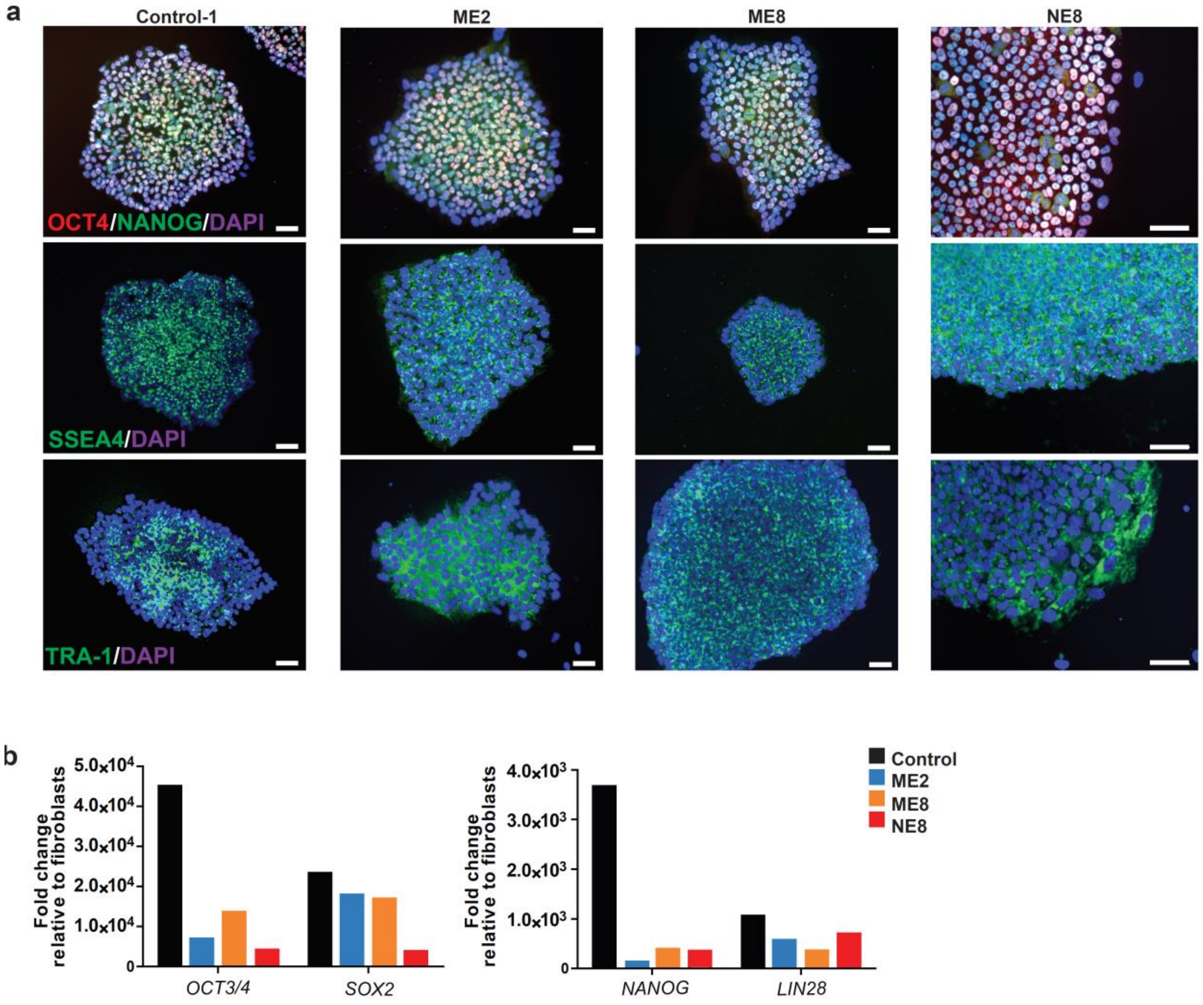


**Supplementary Figure 1: Validation of human induced pluripotent stem cell generation using pluripotency markers** (a) All the lines were positive for the pluripotency markers Oct4, Nanog, SSEA4 and TRA-1 (Scale bar=50 µm). (b) The expression of pluripotent markers increased $10^2$ to $10^5$ fold compared to the average expression in fibroblasts of the same line.

**Supplementary Figure 2**

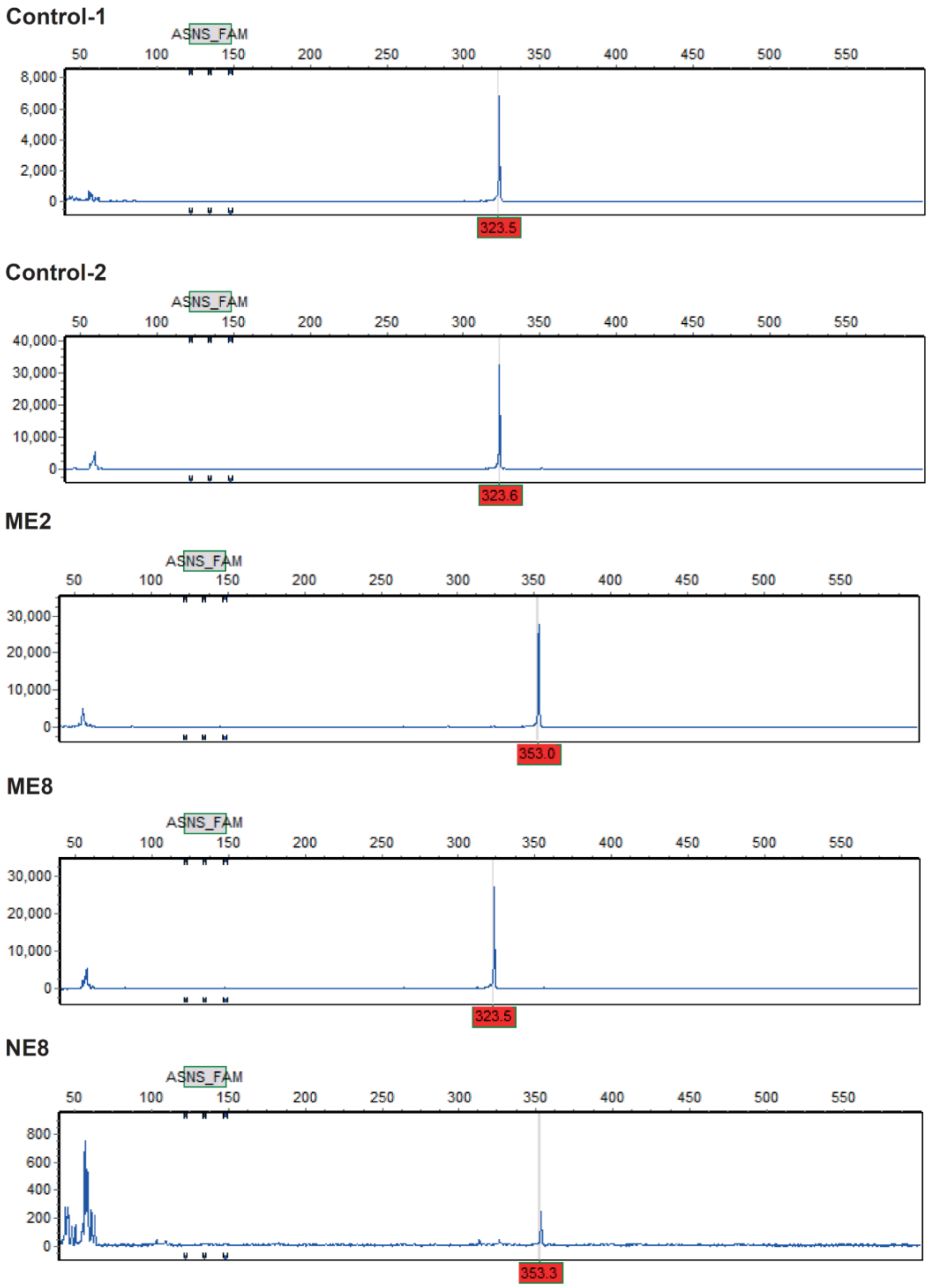


**Supplementary Figure 2: Variable number of tandem repeat (VNTR) polymorphism in the monoamine oxidase A (*MAOA*) promoter** (Corresponding to Table S7). Peaks at position 323 indicate that the individuals carry a 3R allele (*MAOA-L*) whilst peaks at position 353 indicate that the individuals carry a 4R allele (*MAOA-H*).

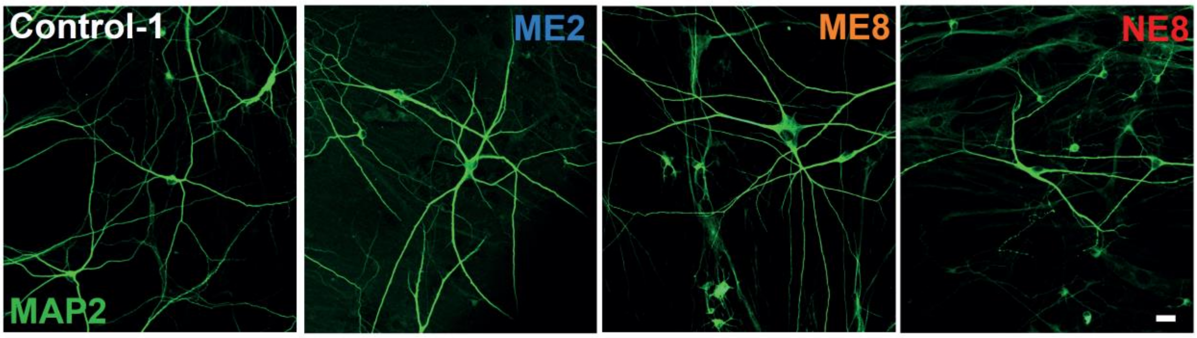


**Supplementary Figure 3: Microtubule Associated Protein 2 (MAP2) staining of DIV 73 DA neurons.** Representative images of DA neurons used in reconstruction and Sholl analysis of neuron dendritic length (Scale bar = 20 µm).

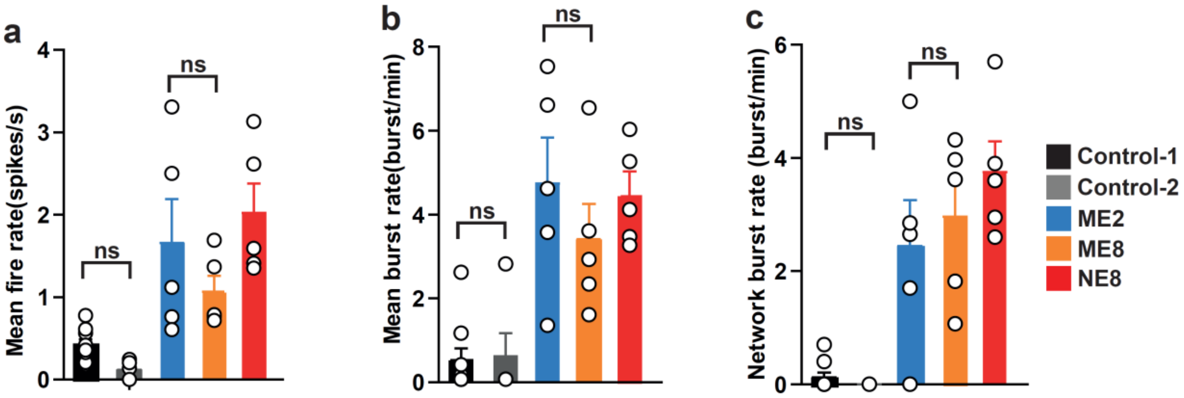


**Supplementary Figure 4: Activity of DIV 73 DA neurons on MEAs separated for all control and patient lines.** No significant differences in activity between control-1 and control-2 and between ME2 and ME8 were observed. All data represent means ± SEM. One-Way ANOVA with Dunnett's correction for multiple testing was used to compare between control and patient lines.

**Supplementary Figure 5**

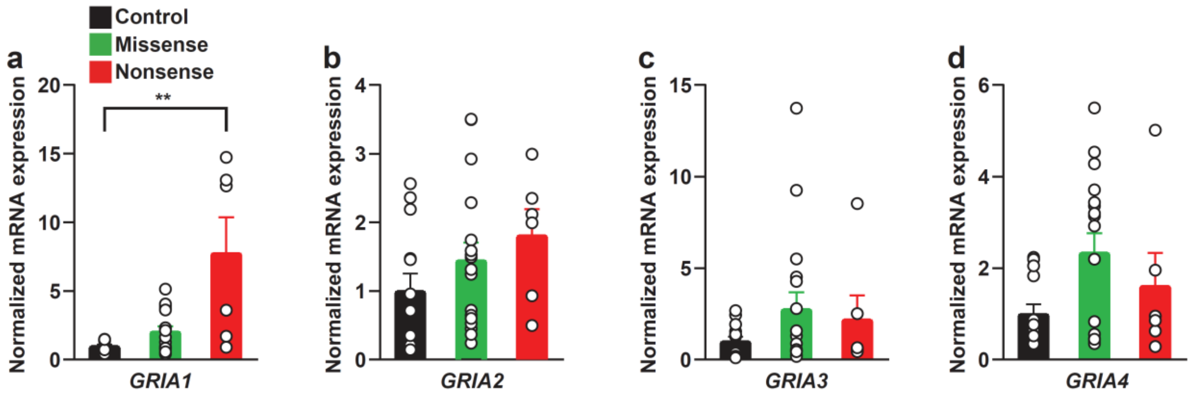


**Supplementary Figure 5: Expression of α-amino-3-hydroxy-5-methyl-4-isoxazolepropionic acid receptor (NMDAR) subunits in control and patient lines.** The expression of genes encoding AMPA receptor subunits *GRIA1* (a, Control vs nonsense *P=0.0077*), *GRIA2* (b), *GRIA3* (c) and *GRIA4* (d) was analyzed by qPCR in DA neurons at day 73 of differentiation. All data represent means ± SEM. One-Way ANOVA with Dunnett's correction for multiple testing was used to compare between control and patient lines. ***p<0.01*. Sample size: N>7 biological replicates across at least 3 batches for all lines.

**Supplementary Figure 6**

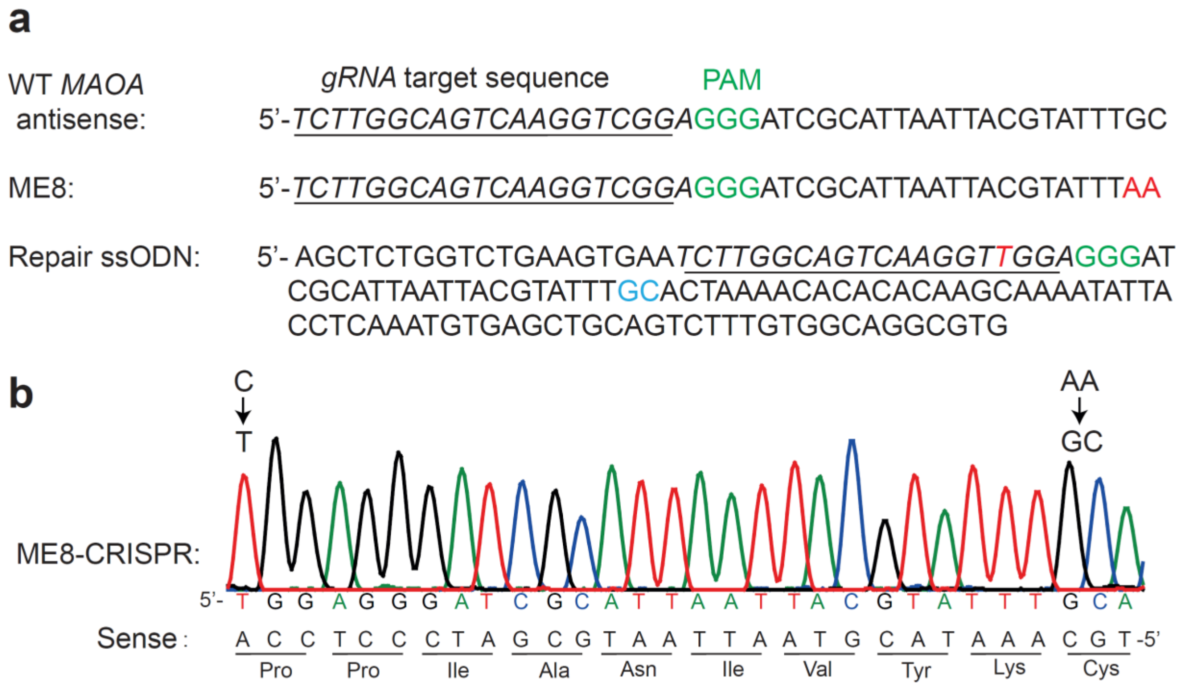


**Supplementary Figure 6: CRISPR/Cas9 mediated correction of the p.C266F (ME8) mutation.** (a) gRNA target sequence (underlined) and protospacer adjustment motif (PAM, marked in green) sequence. The mutation present in patient ME8 (AA, marked in red) was repaired (GC marked in blue) by CRISPR/Cas9 mediated homologous directed recombination (HDR). A synonymous mutation was introduced by substituting a C to T in the repair single-stranded donor oligonucleotides (ssODN) to increase the HDR efficiency (marked in red). (b) The correction of the mutation was confirmed by Sanger sequencing. The predicted top 5 off-target events of CRISPR were ruled out by sequencing the corresponding region (data not shown).

### Supplementary tables

**Supplementary Table S1:** information about the cell lines used in the study, supporting Figure 1

| Cell lines | Gender | Age of biopsy (years) | MAOA Mutation | MAOA VNTR polymorphism (Supplementary Figure 5) | Reference |
|---|---|---|---|---|---|
| Control-1 | Male | 9 | none | 3R, low expression allele (*MAOA-L*) | 3, 4 |
| Control-2 | Male | 30 | none | 3R, low expression allele (*MAOA-L*) | 5 |
| Patient-ME2 | Male | 16 | c.133C>T; p.R45W | 4R, high expression allele (*MAOA-H*) | 8 |
| Patient-ME8 | Male | 10 | c.797_798 delinsTT; p. C266F | 3R, low expression allele (*MAOA-L*) | 7 |
| Patient-NE8 | Male | 21 | c.886C>T; p.Q296* | 4R, high expression allele (*MAOA-H*) | 6 |

**Supplementary Table S2:** Primers for qRT-PCR, supporting Figure 1, 4, 5 and Supplementary Figure 4

| Name | Sequence | References |
|---|---|---|
| B2M Forward | 5’-GCCGTGTGAACCATGTGACT-3’ | 1 |
| B2M Reverse | 5’-GCTTACATGTCTCGATCCCACTT-3’ | |
| MAOA Forward | 5’-CCTTGACTGCCAAGATTCACTTC-3’ | 2 |
| MAOA Reverse | 5’-TGCACTTAATGACAGCTCCCAT-3’ | |
| GRIN1 Forward | 5’-CGCCGCTAACCATAAACAAC-3’ | 3 |
| GRIN1 Reverse | 5’-GGGGAATCTCCTTCTTGACC-3’ | |
| GRIN2A Forward | 5’-AGCTGCTACGGGCAGATG-3’ | |
| GRIN2A Reverse | 5’-CCTGGTAGCCTTCCTCAGTG-3’ | |
| GRIN2B Forward | 5’-GGAGTTCTGGTTCCTACTGGG-3’ | |
| GRIN2B Reverse | 5’-TCTCATGGGAACAGGAATGG-3’ | |
| GRIA1 Forward | 5’-GCAGCAGTGGAAGAATAGTGATG-3’ | |
| GRIA1 Reverse | 5’-ATCACCTTCACCCCATCGTA-3’ | |
| GRIA2 Forward | 5’-GCTTGGTGCTAAATTGCTGT-3’ | |
| GRIA2 Reverse | 5’-TCCAAGAAAAGTAGAGCATCCA-3’ | |
| GRIA3 Forward | 5’-TTCCCACTGGAGGCATGTG-3’ | |
| GRIA3 Reverse | 5’-CATCAGCAATATTCGTGTCATGC-3’ | |
| GRIA4 Forward | 5’-TGCTGCAACTAAGACCTTCGTTAC-3’ | |
| GRIA4 Reverse | 5’-TCGAGTATCCCCTGTCTGTGTC-3’ | |

**Supplementary data tables:**

**Supplementary Table S3: statistics related to Figure 1 and Supplementary Figure 2**.

| **Panel** | | LINE | VALUE | SEM | P-VALUE | COMPARED TO |
|---|---|---|---|---|---|---|
| **Fig. 1e** | $TH^+$/$MAP2^+$ neurons | Control-1 | 94.9 | 1.05 | | |
| | | Control-2 | 86.9 | 3.09 | | |
| | | ME2 | 92.2 | 2.34 | | |
| | | ME8 | 88.8 | 2.58 | | |
| | | NE8 | 91.12 | 1.97 | | |
| **Fig. 1f** | Normalized MAOA mRNA expression (to control-1) | Control-1 | 1.00 | 0.12 | | |
| | | Control-2 | 0.70 | 0.24 | | |
| **Fig. 1g** | Normalized MAOA mRNA expression (to control pooled) | Control | 1.00 | 0.18 | | |
| | | ME2 | 0.84 | 0.11 | | |
| | | ME8 | 0.70 | 0.13 | | |
| | | NE8 | 0.32 | 0.08 | *P=0.0076* | Control |
| **Sup. Fig 1b** | OCT3/4 fold change compared to fibroblast | Control | $4.5*10^4$ | | | |
| | | ME2 | $0.68*10^4$ | | | |
| | | ME8 | $1.35x10^4$ | | | |
| | | NE8 | $0.39*10^4$ | | | |
| **Sup. Fig 1b** | SOX2 fold change compared to fibroblast | Control | $2.32*10^4$ | | | |
| | | ME2 | $1.76*10^4$ | | | |
| | | ME8 | $1.68*10^4$ | | | |
| | | NE8 | $0.34*10^4$ | | | |
| **Sup. Fig 1b** | NANOG fold change compared to fibroblast | Control | $3.66*10^3$ | | | |
| | | ME2 | $0.13*10^3$ | | | |
| | | ME8 | $0.38*10^3$ | | | |
| | | NE8 | $0.32*10^3$ | | | |
| **Sup. Fig 1b** | LIN28 fold change compared to fibroblast | Control | | | | |
| | | ME2 | $0.57*10^3$ | | | |
| | | ME8 | $0.35*10^3$ | | | |
| | | NE8 | $0.69*10^3$ | | | |

Sample size for the MAOA mRNA expression analysis control-1 N=13, control 2 N=7, ME2 N=12, ME8 N=12, NE8 N=12. N=number of replicates across as least 3 individual differentiations. Sample size for the calculation of TH positive neurons control-1 N=15, control-2 N=16, ME2 N=15, ME8 N=15, NE8 N=15. N=number of technical replicates across at least 3 individual differentiations. All data represent means ± SEM. One-Way ANOVA with Dunnett's correction for multiple testing was used to compare between patient lines and control lines. The pluripotent stem cell makers were tested by qPCR at Stem Cell Technology Center and passed their criteria for hiPSCs.

**Supplementary Table S4: statistics related to Figure 2.**

| **Panel** | | LINE | VALUE | SEM | P-VALUE | COMPARED TO |
|---|---|---|---|---|---|---|
| **Fig. 2b** | Soma area (µ$m^2$) | Control-1 | 237.33 | 22.1 | | |
| | | ME2 | 254.16 | 17.8 | | |
| | | ME8 | 470.70 | 50.7 | *P<0.001* | Control-1 |
| | | NE8 | 271.28 | 31.0 | | |
| **Fig. 2c** | Dendritic nodes (n) | Control-1 | 10.3 | 1.08 | | |
| | | ME2 | 15.3 | 1.39 | | |
| | | ME8 | 32.4 | 2.88 | *P<0.001* | Control-1 |
| | | NE8 | 14.3 | 1.18 | | |
| **Fig. 2d** | Dendritic length (µm) | Control | 1430 | 88.5 | | |
| | | ME2 | 1913 | 89.5 | | |
| | | ME8 | 3723 | 213.3 | *P<0.001* | Control-1 |
| | | NE8 | 1685 | 96.1 | | |
| **Fig. 2e** | Primary dendrites (n) | Control | 4.5 | 0.30 | | |
| | | ME2 | 4.3 | 0.33 | | |
| | | ME8 | 5.4 | 0.29 | | |
| | | NE8 | 4.3 | 0.38 | | |
| **Fig. 2g** | Convex hull: Covered surface (µ$m^2$) | Control | 70953 | 5443 | | |
| | | ME2 | 122746 | 12394 | *P<0.05* | |
| | | ME8 | 214870 | 14907 | *P<0.001* | Control-1 |
| | | NE8 | 90008 | 5898 | | |
| **Fig. 2h** | Synapse density (n/10µm) | Control | 1.749 | 0.087 | | |
| | | ME2 | 1.178 | 0.0828 | *P=0.0001* | |
| | | ME8 | 1.177 | 0.056 | *P=0.0001* | |
| | | NE8 | 1.406 | 0.079 | *P=0.0054* | Control1 |

Sample size for the morphological reconstruction of the somatodendritic compartment N=20 neurons from 3 neuronal preparations for all lines. Sample size for the synapse density: control-1 N=25, ME2 N=23, ME8 N=25, NE8 N=26 neurons from 3 neuronal preparations for all lines. All data represent means ± SEM. One-Way ANOVA with Dunnett's correction for multiple testing was used to compare between patient lines and the control-1 line.

**Supplementary Table S5: statistics related to Figure 3 and Supplementary Figure 4**

| Panel | | LINE | VALUE | SEM | P-VALUE | COMPARED TO |
|---|---|---|---|---|---|---|
| **Fig. 3d** | Firing rate (spikes/s) | Control-1 | 0.31 | 0.06 | | |
| | | Missense | 1.33 | 0.28 | *P<0.001* | Control-1 |
| | | Nonsense | 1.99 | 0.36 | *P<0.001* | Control-1 |
| **Fig. 3e** | Mean burst rate (N/min) | Control-1 | 0.49 | 0.26 | | |
| | | Missense | 4.02 | 0.69 | *P<0.001* | Control-1 |
| | | Nonsense | 4.42 | 0.53 | *P<0.001* | Control-1 |
| **Fig. 3f** | Network burst rate (N/min) | Control | 0.08 | 0.06 | | |
| | | Missense | 2.69 | 0.50 | *P<0.001* | Control-1 |
| | | Nonsense | 3.75 | 0.53 | *P<0.001* | Control-1 |
| **Sup. Fig. 4a** | Firing rate (spikes/s) | Control-1 | 0.42 | 0.06 | | |
| | | Control-2 | 0.19 | 0.03 | | |
| | | ME2 | 1.62 | 0.36 | | |
| | | ME8 | 1.06 | 0.20 | | |
| | | NE8 | 1.99 | 0.53 | | |
| **Sup. Fig. 4b** | Mean burst rate (N/min) | Control-1 | 0.46 | 0.29 | | |
| | | Control-2 | 0.55 | 0.55 | | |
| | | ME2 | 4.67 | 1.10 | | |
| | | ME8 | 3.37 | 0.85 | | |
| | | NE8 | 4.42 | 0.53 | | |
| **Sup. Fig. 4c** | Network burst rate (N/min) | Control-1 | 0.12 | 0.09 | | |
| | | Control-2 | 0 | 0 | | |
| | | ME2 | 2.44 | 0.81 | | |
| | | ME8 | 2.94 | 0.64 | | |
| | | NE8 | 3.75 | 0.54 | | |

Sample size for MEA recording of neuronal network activity: control-1 N=9, control-2 N=5, ME2 N=5, ME8 N=5, NE8 N=5 – recordings from at least 3 different neuronal preparations for all lines. All data represent means ± SEM. One-Way ANOVA with Dunnett's correction for multiple testing was used to compare between patient lines and control lines.

**Supplementary Table S6: statistics related to Figure 4.**

| Panel | | LINE | VALUE | SEM | P-VALUE | COMPARED TO |
|---|---|---|---|---|---|---|
| **Fig. 4b** | Capacitance (pF) | Control-1 | 29.4 | 2.21 | | |
| | | Missense | 28.7 | 2.06 | | |
| | | Nonsense | 34.7 | 4.25 | | |
| **Fig. 4c** | Membrane resistance (MΩ) | Control-2 | 1476 | 227.6 | | |
| | | Missense | 1392 | 130.6 | | |
| | | Nonsense | 1218 | 242.3 | | |
| **Fig. 4d** | Vrmp (MV) | Control | -34.7 | 2.40 | | |
| | | Missense | -29.6 | 2.71 | | |
| | | Nonsense | -34.6 | 2.62 | | |
| **Fig. 4e** | Rheobase (pA) | Control | 22.0 | 2.31 | | |
| | | Missense | 27.2 | 4.41 | | |
| | | Nonsense | 25.0 | 4.37 | | |
| **Fig. 4f** | AP threshold (mV) | Control | -27.9 | 1.21 | | |
| | | Missense | -28.1 | 0.93 | | |
| | | Nonsense | -30.6 | 1.75 | | |
| **Fig. 4g** | AP amplitude (mV) | Control | 97.9 | 2.66 | | |
| | | Missense | 92.1 | 3.70 | | |
| | | Nonsense | 90.0 | 8.03 | | |
| **Fig. 4i** | sPSC amplitude (pA) | Control-1 | 22.0 | 2.42 | | |
| | | Missense | 19.2 | 1.93 | | |
| | | Nonsense | 16.5 | 1.34 | | |
| **Fig. 4k** | sPSC frequency (Hz) | Control-2 | 1.45 | 0.22 | | |
| | | Missense | 1.00 | 0.21 | | |
| | | Nonsense | 0.83 | 0.28 | | |
| **Fig. 4l** | *GRIN1* expression (normalized) | Control | 0.88 | 0.14 | | |
| | | Missense | 1.11 | 0.23 | | |
| | | Nonsense | 1.20 | 0.39 | | |
| **Fig. 4m** | *GRIN2A* expression (normalized) | Control | 0.90 | 0.12 | | |
| | | Missense | 1.85 | 0.42 | *P=0.0363* | Control |
| | | Nonsense | 2.34 | 0.60 | *P=0.0384* | Control |
| **Fig. 4n** | *GRIN2B* expression (normalized) | Control | 0.94 | 0.17 | | |
| | | Missense | 1.79 | 1.76 | *P=0.0169* | Control |
| | | Nonsense | 2.99 | 0.49 | *P=0.0002* | Control |
| **Fig. 4p** | NMDA current area under curve (pA*sec) | Control | -145767 | 30152 | | |
| | | Missense | -290974 | 71569 | *P=0.0287* | Control |
| **Sup. Fig. 5a** | *GRIA1* expression (normalized) | Control | 1.00 | 0.11 | | |
| | | Missense | 2.05 | 0.37 | | |
| | | Nonsense | 7.77 | 2.59 | | |
| **Sup. Fig.5b** | *GRIA2* expression (normalized) | Control | 1.00 | 0.25 | | |
| | | Missense | 1.45 | 0.26 | | |
| | | Nonsense | 1.81 | 0.38 | | |
| **Sup. Fig. 5c** | *GRIA3* expression (normalized) | Control | 1.00 | 0.26 | | |
| | | Missense | 2.76 | 0.91 | | |
| | | Nonsense | 2.21 | 1.30 | | |
| **Sup. Fig. 5d** | *GRIA4* expression (normalized) | Control | 1.00 | 0.21 | | |
| | | Missense | 2.34 | 0.42 | | |
| | | Nonsense | 1.61 | 0.71 | | |

Control-1 and control-2 and missense line ME2 and ME8 are pooled in all analyses. Sample size for single cell recording of intrinsic properties: control N=27, missense N=21, nonsense N=12. Sample size for recording of spontaneous activity: control N=19, missense N=16, nonsense N=10. Sample size for NMDAR subunit mRNA expression analysis: control N>25, missense N>26, nonsense N>13. Sample size for NMDA puff: control N=16, missense N=13, nonsense N=4. Sample size for AMPAR subunit mRNA expression analysis: control N=13, missense N=17, monsense N=6. Data are from at least 2 different neuronal preparations for all lines. All data represent means ± SEM. One-Way ANOVA with Dunnett's correction for multiple testing was used to compare between patient lines and control lines.

**Supplementary Table S7: statistics related to Figure 5 and Supplementary Figure 6.**

| **Panel** | | LINE | VALUE | SEM | P-VALUE | COMPARED TO |
|---|---|---|---|---|---|---|
| **Fig. 5b** | *GRIN1* expression (normalized) | Control | 0.88 | 0.14 | | |
| | | ME8 | 1.04 | 0.26 | | |
| | | ME8-CRISPR | 1.33 | 0.31 | | |
| **Fig. 5c** | *GRIN2A* expression (normalized) | Control | 0.90 | 0.12 | | |
| | | ME8 | 1.82 | 0.43 | *P=0.0217* | Control |
| | | ME8-CRISPR | 0.85 | 0.22 | | |
| **Fig. 5d** | *GRIN2B* expression (normalized) | Control | 0.94 | 0.17 | | |
| | | ME8 | 1.63 | 0.34 | *P=0.0186* | Control |
| | | ME8-CRISPR | 0.88 | 0.25 | | |
| **Fig. 5f** | NMDA current area under curve (pA*sec) | Control | -145767 | 30152 | | |
| | | ME8 | -302399 | 83274 | *P=0.0256* | Control |
| | | ME8-CRISPR | -75377 | 20782 | *P=0.0180* | ME8 |
| **Fig. 5h** | Firing rate (spikes/s) | Control | 0.76 | 0.09 | | |
| | | ME8 | 1.51 | 0.23 | *P=0.00043* | Control |
| | | ME8-CRISPR | 0.59 | 0.08 | *P=0.00011* | ME8 |
| **Fig. 5i** | Mean burst rate (N/min) | Control | 3.23 | 0.52 | | |
| | | ME8 | 5.11 | 0.81 | *P=0.0046* | Control |
| | | ME8-CRISPR | 2.16 | 0.62 | *P=0.0005* | ME8 |

Control-1 and control-2 are pooled in all analyses. Sample size for NMDAR subunit mRNA expression analysis: control N>25, ME8 N>14, ME8-CRISPR N>7. Data are from at least 3 different neuronal preparations for all lines Sample size for NMDA puff: Control N=16, ME8 N=11, ME8-CRISPR N=8. Data are from at least 2 different neuronal preparations for all lines. Sample size for MEA recording of neuronal network activity: control N=23, ME8 N=12, ME8-CRISPR N=13. Sample size for developmental timeline of MEA activity: control N=9, ME8 N=9, ME8-CRISPR N=9. Data are from at least 2 different neuronal preparations for all lines. Data represent means ± SEM. One-Way ANOVA with Dunnett's correction for multiple testing was used to compare control, ME8 and ME8-CRISPR lines.
